# Dopamine Activates Astrocytes in Prefrontal Cortex via α1-Adrenergic Receptors

**DOI:** 10.1101/2022.07.19.500710

**Authors:** Silvia Pittolo, Sae Yokoyama, Drew D. Willoughby, Charlotte R. Taylor, Michael E. Reitman, Vincent Tse, Zhaofa Wu, Roberto Etchenique, Yulong Li, Kira E. Poskanzer

**Affiliations:** Department of Biochemistry & Biophysics, University of California, San Francisco, San Francisco, CA, USA; Max Delbrück Center for Molecular Medicine in the Helmholtz Association, Robert-Rössle-Str. 10, 13125 Berlin, Germany; Neuroscience Graduate Program, University of California, San Francisco, San Francisco, CA, USA; State Key Laboratory of Membrane Biology, Peking University School of Life Sciences, Beijing 100871, China; Peking-Tsinghua Center for Life Sciences, Peking University, Beijing 100871, China; Departamento de Química Inorgánica, Analítica y Química Física, INQUIMAE, Facultad de Ciencias Exactas y Naturales, Universidad de Buenos Aires, CONICET, Intendente Güiraldes 2160, Ciudad Universitaria, Pabellón 2, Buenos Aires (C1428EGA), Argentina; Kavli Institute for Fundamental Neuroscience, San Francisco, CA, USA

**Keywords:** Prefrontal cortex, astrocytes, dopamine, adrenergic receptors, calcium, two-photon imaging, fiber photometry, cAMP, aversive stimulus, ATP, receptor interactions, GPCR signaling

## Abstract

The prefrontal cortex (PFC) is a hub for cognitive control, and dopamine profoundly influences its functions. In other brain regions, astrocytes sense diverse neurotransmitters and neuromodulators and, in turn, orchestrate regulation of neuroactive substances. However, basic physiology of PFC astrocytes, including which neuromodulatory signals they respond to and how they contribute to PFC function, is lacking. Here, we characterize divergent signaling signatures in astrocytes of PFC and primary sensory cortex in mice, which are linked to differential responsivity to locomotion. We find that PFC astrocytes express receptors for dopamine, but are unresponsive through the G_s_/G_i_-cAMP pathway. Instead, fast calcium signals in PFC astrocytes are time-locked to dopamine release, and are mediated by α1-adrenergic receptors both *ex vivo* and *in vivo*. Further, we describe dopamine-triggered regulation of extracellular ATP at PFC astrocyte territories. Thus, we identify astrocytes as active players in dopaminergic signaling in PFC, contributing to PFC function though neuromodulator receptor crosstalk.

**Graphical Abstract:** 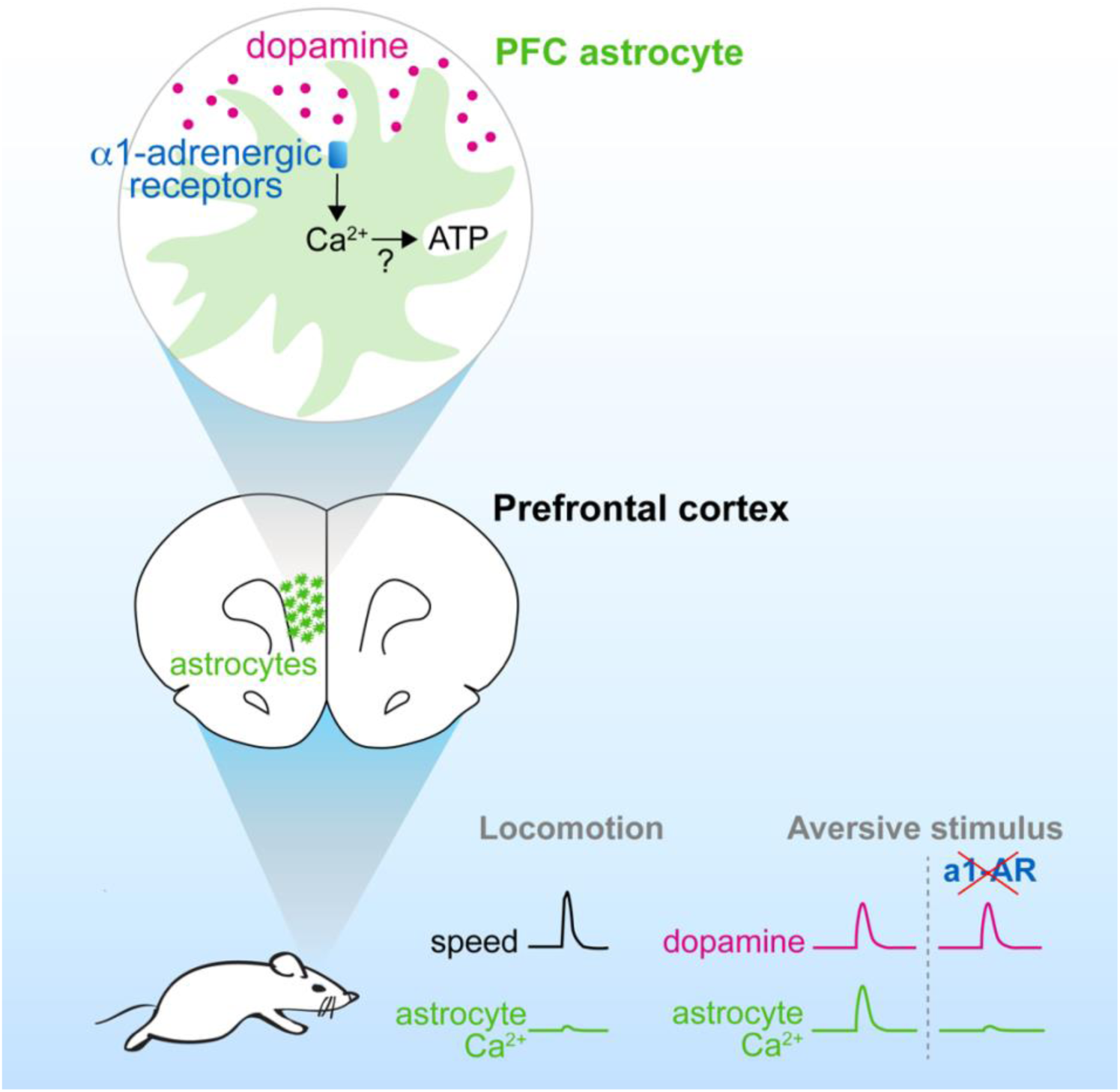

## Introduction

The prefrontal cortex (PFC) is a higher order association cortex that integrates and holds sensory and cognitive information from other brain areas to organize and execute behavior (Fuster et al., 2000). The PFC is involved in fundamental and diverse processes, including working memory and attention (Funahashi et al., 1989; Fuster and Alexander, 1971; Kesner et al., 1996), behavioral flexibility and planning (Dias et al., 1996; Ragozzino et al., 1999), and sustained processing of stress, fear and emotions (George et al., 1995; Hariri et al., 2003; Kim et al., 2003; Milad and Quirk, 2002). Despite its importance, how the PFC performs many aspects of its complex functions is still poorly understood. For instance, whether persistent activity of individual PFC neurons or rather network dynamics underlie the ability of the PFC to hold information over multi-second delays during working memory tasks is subject of current debate (Barbosa et al., 2020; Cavanagh et al., 2018; Constantinidis et al., 2018; Inagaki et al., 2019; Park et al., 2019; Spaak et al., 2017).

While prefrontal circuits are fundamental for the top-down control of behavior, ascending arousal systems—including the mesocortical dopamine (DA) pathway—are so necessary for PFC executive functions that their disruption recapitulates PFC lesions (Brozoski et al., 1979). Dopaminergic projections to the PFC are particularly sensitive to stressful and aversive stimuli (Abercrombie et al., 1989; Lammel et al., 2012; Thierry et al., 1976; Vander Weele et al., 2018). However, how both fast (phasic) and prolonged (tonic) temporal patterns of DA release play specific roles in PFC computations is unclear (Lohani et al., 2019), with evidence for bidirectional or opposing effects on the excitability of prefrontal neuron subtypes (Anastasiades et al., 2019; Chen et al., 2007; Gao and Goldman-Rakic, 2003; Gao et al., 2003; Huang et al., 2004; Kröner et al., 2007; Matsuda et al., 2006; Seamans et al., 2001; Vijayraghavan et al., 2007) which ultimately contribute to the complex patterns of sustained and dynamic circuit activity underlying PFC functions.

Astrocytes—the most abundant non-neuronal brain cell type—are also well positioned to receive and process neuronal signals, as they are equipped with an array of receptors for neurotransmitters and neuromodulators (Porter and McCarthy, 1997) and have wide, non-overlapping territories, each encompassing thousands of synapses (Bushong et al., 2002). Many lines of research show that astrocytes are in a bidirectional dialogue with neurons, by sensing neuronal activity through G protein-coupled receptors (GPCRs) (Kofuji and Araque, 2021), internally computing through Ca^2+^ and cAMP signals (Oe et al., 2020; Srinivasan et al., 2016), and regulating neuroactive substances such as glutamate (Bezzi et al., 2004; Yang et al., 2019) and ATP (Cao et al., 2013; Pascual et al., 2005; Zhang et al., 2003) that can influence local synaptic plasticity and network connectivity on multiple spatiotemporal scales (Panatier et al., 2011; Perea and Araque, 2007; Poskanzer and Yuste, 2016). Previous work shows that chemical ablation of PFC astrocytes, blockade of glutamate/glutamine homeostasis or DA uptake and metabolism, impaired gliotransmission, Ca^2+^ or ATP signals, or lack of GABA_B_ receptors in PFC astrocytes cause depressive (Banasr and Duman, 2008; Cao et al., 2013; John et al., 2012; Lee et al., 2013) or autism-like behaviors (Wang et al., 2021), and interfere with working memory (Lima et al., 2014; Mederos et al., 2021; Petrelli et al., 2020; Sardinha et al., 2017). With these notable exceptions, decades of research have largely neglected the contribution of astrocyte physiology to PFC regulation.

Here, we use a combination of *in vivo* two-photon (2P) imaging, fiber photometry, and *ex vivo* imaging of calcium (Ca^2+^), cAMP, neuromodulators, and ATP to explore astrocyte signals in PFC. We first characterize endogenous Ca^2+^ dynamics of PFC astrocytes *in vivo* through GRIN lens imaging (Levene et al., 2004), and compare them to primary sensory cortical astrocytes. We find that PFC astrocytes display unique spatiotemporal signals compared to those in primary visual cortex (V1), and lack responsiveness to locomotory activity observed in sensory cortex (Paukert et al., 2014; Wang et al., 2019). We next demonstrate that PFC astrocytes express dopamine receptors (DR), but signal through fast, sustained Ca^2+^ mobilizations rather than the canonical G_s_/G_i_-cAMP pathway. Unexpectedly, we find that DA in PFC elicits astrocyte activation through the G_q_-coupled α1-AR, which we confirmed by pharmacological screening with DR and adrenergic receptor (AR) inhibitors both in acute slices and *in vivo*. Finally, we show that PFC astrocytes can regulate extracellular ATP in response to DA, which may contribute to neuronal computation in PFC. Together, our data demonstrate that astrocytes in PFC sense neuromodulators and behavioral stimuli differently than sensory cortical astrocytes. By exploring the physiology of PFC astrocytes, we uncover functional crosstalk—both *ex vivo* and *in vivo*—between DA and receptors for norepinephrine (NE).

## Results

### PFC astrocytes exhibit more single-cell restricted Ca^2+^ activity compared to sensory cortex

Since PFC is an associative cortical area (Fuster et al., 2000), we hypothesized that astrocyte Ca^2+^ in PFC may have unique properties compared to primary sensory cortices, where population-level bursts of activity have been well documented (Bekar et al., 2008; Ding et al., 2013; Slezak et al., 2019; Srinivasan et al., 2016; Wang et al., 2019). To test this, we compared spontaneous astrocyte Ca^2+^ activity in PFC and primary visual cortex (V1) using two-photon (2P) microscopy in head-fixed mice. To gain imaging access, we implanted either a GRIN lens (Levene et al., 2004) over PFC or a cranial window over V1 in mice expressing Lck-GCaMP6f (Fig 1A) (Shigetomi et al., 2010; Srinivasan et al., 2016) under the astrocyte-specific promoter *GfaABC_1_D*. GRIN lens positioning was confirmed post-mortem (Fig S1A), and GFAP staining was used to assess low astrocyte reactivity around the implant (Fig S1B).

**Figure 1:**
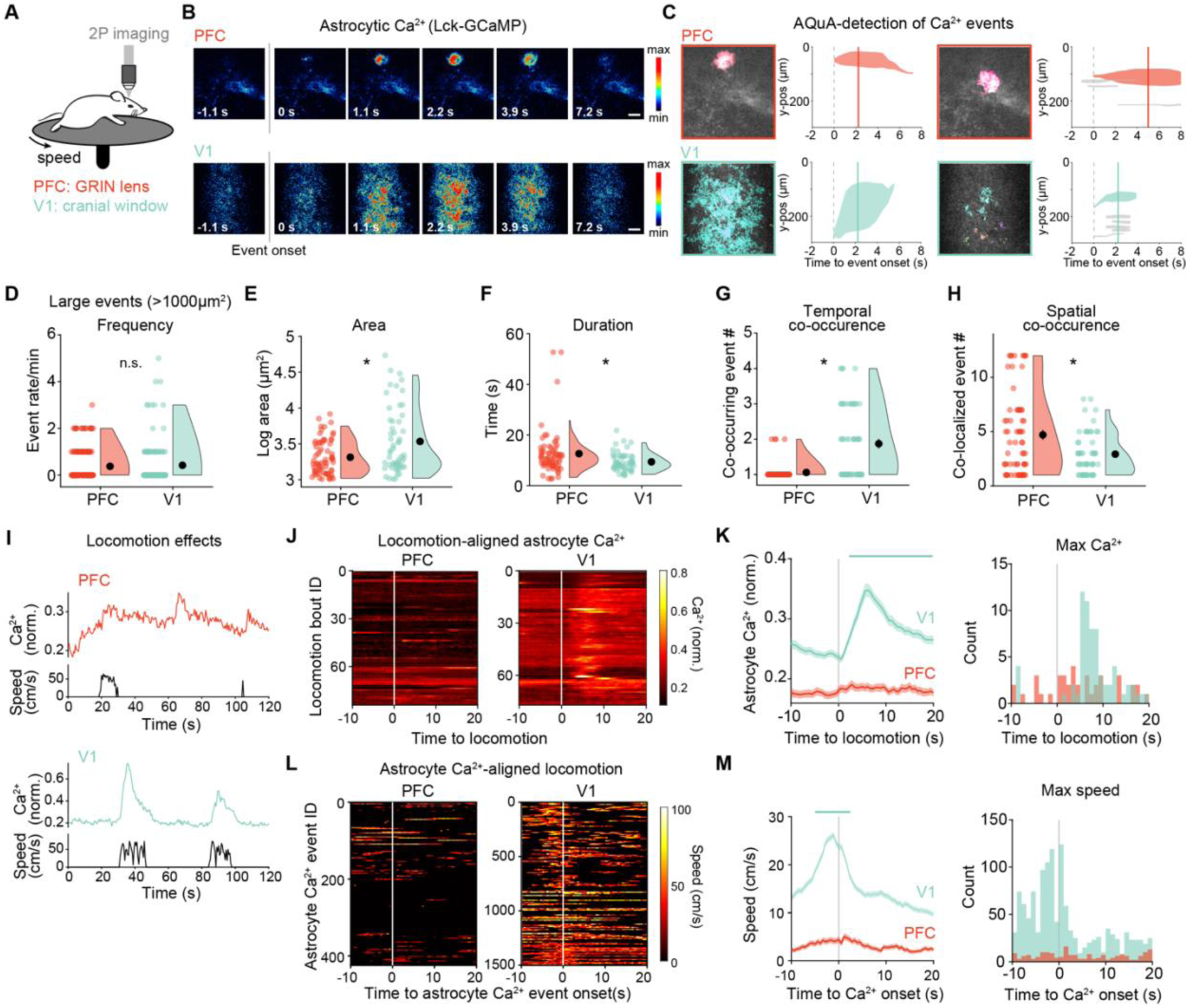
PFC astrocytes exhibit region-specific Ca^2+^ activity. (A) Experimental setup for head-fixed 2P imaging of astrocyte Ca^2+^ in PFC (via GRIN lens) or V1 (cranial window) in vivo. (B) Representative frames of pseudocolored astrocytic GCaMP6f fluorescence in PFC (top) or V1 (bottom), relative to Ca^2+^ event onset. Scale bars = 50 µm. (C) Two examples of large AQuA-detected Ca^2+^-events each in PFC (red, top) and V1 (green, bottom). Fields-of-view = 300 x 300 µm^2^. To the right of each image is the corresponding timecourse of all detected events within 10 s, with the time of onset of the largest event at t = 0 and the solid line indicating the frame displayed at left. Events <1000 µm^2^ are in gray. (D–H) Large (>1000 µm^2^) astrocyte Ca^2+^-event features vary between brain regions. Events occur at similar rates in PFC and V1 (D), but in PFC are (E) smaller and (F) longer than in V1. Events in PFC (G) co-occur with other events less than in V1, but (H) tend to repeat more at the same spatial location. Data shown as all bins/events (colored dots), 5^th^–95^th^ percentile distribution (violins), and mean ± sem (black dots and error bars). Event rate (min^-1^): 0.38±0.05 (PFC), 0.42±0.08 (V1); area (µm^2^): 2422±190 (PFC), 6639±1346 (V1); duration (s): 12.6±1.1 s (PFC), 9.4±0.5 s (V1); temporal co-occurrence: 1.06±0.03 (PFC), 1.87±0.13 (V1); spatial co-occurrence: 4.7±0.5 (PFC), 2.9±0.3 (V1). Wilcoxon rank-sum test; *, p < 0.05; p = 0.6288 (frequency), p = 0.0122 (area), p = 0.0116 (duration), p < 10^-4^ (co-occurring), p = 0.0340 (co-localized). PFC: n = 180 60-s bins, 68 events, 4 mice; V1: n = 130 60-s bins, 55 events, 3 mice. (I–M) Locomotion does not induce population-wide astrocyte Ca^2+^ in PFC. (I) Example time course of normalized astrocyte Ca^2+^ (colored trace, top) and corresponding mouse speed (black, bottom) in PFC and VI. (J–K) Astrocyte Ca^2+^ traces aligned to the onset of locomotion (t = 0), shown as heatmaps for all recordings (J), average traces ± sem (K, left) and binned distribution of maximum Ca^2+^ change (K, right). In (K), line above mean traces indicates significant change from average Ca^2+^ at t < 0. Shuffle test, 10000 pair-wise shuffles; p < 0.01, Bonferroni correction for multiple comparisons. PFC: n = 84 bouts, 4 mice; V1: n = 77 bouts, 3 mice. (L–M) Animal speed aligned to onset of astrocyte Ca^2+^ events (t = 0), shown as heat map for all recordings (L), average traces ± sem (M, left), and binned distribution of maximal speed (M, right). In (M), line above mean traces indicates significant increase above the average speed for the entire window. Shuffle test, 10000 pair-wise shuffles; p < 0.01, Bonferroni correction for multiple comparisons. PFC: n = 424 events, 4 mice; V1: n = 1501 events, 3 mice.

Using event-based image analyses to detect the area of astrocyte Ca^2+^ independent of pre-determined regions-of-interest (Wang et al., 2019), we observed that the largest astrocyte Ca^2+^ signals in PFC often appeared to be the size of a single astrocyte (∼50×50 µm; Fig 1B–C top), whereas the population-level, burst-like events in V1 span the entire imaging field (here: 300×300 µm; Fig 1B–C bottom), as previously reported (Slezak et al., 2019; Srinivasan et al., 2016; Wang et al., 2019). We thus focused on these larger events (>1000 µm^2^) for quantitative comparison (Fig 1D–H), and found that while astrocyte Ca^2+^ events occur at the same rate in PFC and V1 (Fig 1D), they are smaller (Fig 1E) and last longer (Fig 1F) in PFC. When examining the network properties of astrocyte Ca^2+^, we found that events in PFC are less synchronous with other PFC events (Fig 1G), but tend to repeat more at the same locations in the imaging field compared to those in V1 (Fig 1H). Although less obvious by eye, smaller Ca^2+^ events (<1000 µm^2^) also differ between PFC and VI (Fig S1C–G), indicating that overall Ca^2+^ dynamics may be driven by different mechanisms depending on brain region. Together, these data demonstrate that astrocyte Ca^2+^ dynamics in PFC are significantly different from those in V1, suggesting that astrocytes in PFC may play different functional roles than in primary sensory cortex.

### Population-level astrocyte Ca^2+^ activity in PFC is not tightly linked to locomotion

Since burst-like astrocyte activity in V1 is known to be driven by locomotion (Paukert et al., 2014; Slezak et al., 2019; Wang et al., 2019), we next wondered whether the observed differences in astrocyte Ca^2+^ dynamics in PFC compared to V1 were due to differences in astrocytic response to locomotion (Movie 1). To examine this, we aligned population-level astrocyte Ca^2+^ traces to the onset of locomotion bouts (Fig 1J–K and Fig S1H–I). Similar to previous results (Paukert et al., 2014; Slezak et al., 2019), average astrocyte Ca^2+^ in V1 significantly increases soon after locomotion onset (2.2 s), peaks at 6.2 s, and stays above baseline for the remaining 15 s (Fig 1K left, green). When we plot the distribution of the times at which we detect the maximum change in astrocyte Ca^2+^ (Fig 1K, right), we observe a clear peak 6–9 s after locomotion onset. In contrast, in PFC astrocytes we did not find significant and sustained Ca^2+^ increases at the onset of locomotion on average, and no clear peak for maximum change across trials is evident (Fig 1K, red). These results indicate that whereas V1 astrocytes are recruited soon after an animal begins locomotion, PFC astrocytes are not activated by locomotion on average, although we do not exclude the possibility that a few astrocytes or astrocytic domains are locomotion-linked.

These results suggest that astrocytic Ca^2+^ is not linked to locomotion in PFC. We next wondered whether Ca^2+^ events in PFC astrocytes are instead involved in the generation of locomotion . To explore this, we aligned locomotion traces to the onset of astrocyte Ca^2+^ events (Fig 1L–M, Fig S1J–K), and found no times around Ca^2+^ onsets when animal speed significantly deviated from average (Fig 1M left, red). When mice did move, the maximum speed was equally distributed over the time window around astrocyte Ca^2+^ (Fig 1M, right), suggesting that astrocytic Ca^2+^ activity in PFC is unlinked from locomotion. In contrast to PFC, astrocyte Ca^2+^-aligned locomotion analysis in V1 shows that speed increases starting at -5.1 s before Ca^2+^ event onset and until 2.2 s after, and peaks -1.1 s before Ca^2+^ onset (Fig 1M left, green), which is in accordance with previous observations that locomotion initiates Ca^2+^ activity in V1 astrocytes (Paukert et al., 2014; Slezak et al., 2019) (Fig 1J–K). Together, these results indicate that astrocyte activity in PFC differs significantly from that observed in V1, both in terms of Ca^2+^ event dynamics and their relationship with locomotion.

### PFC astrocytes express dopamine receptors

Because it has been previously shown that burst-like astrocyte population dynamics are mediated by noradrenergic signaling (Bekar et al., 2008; Ding et al., 2013; Paukert et al., 2014) and PFC astrocytes do not display this bursting activity (Fig 1B–C), we wondered whether another neuromodulator may be involved in the astrocyte activity we observe in PFC *in vivo*. We reasoned that since dopamine (DA) is an important neuromodulatory input for PFC neurons (Brozoski et al., 1979; Thierry et al., 1976), DA may also activate PFC astrocytes. To explore this possibility, we first examined DA receptor expression in prefrontal astrocytes by crossing the transgenic reporter lines *Drd1a*-tdTomato (Shuen et al., 2008) or *Drd2*-EGFP (Gong et al., 2003), which have been widely used to identify expression of D1R or D2R in neurons, to the astrocyte-specific reporter lines *Aldh1l1*-EGFP (Tsai et al., 2012) or *Aldh1l1*-tdTomato (Gong et al., 2003) (Fig 2A). We immunostained for the fluorophores in young adult mice (P29–32) and determined colocalization of these reporters in cell somata across PFC layers (Fig 2C–D and Fig S2A). We found that 13±1% of all Aldh1l1^+^ cells colocalize with D1R and 14±1% colocalize with D2R (Fig 2C). Conversely, 18±2% of all D1R^+^ cells, and 41±3% of all D2R^+^ cells are Aldh1l1^+^ (Fig 2D). For both receptors, colocalization with Aldh1l1 in astrocytes was maximal in layer 1, consistent with the prevalence of neuronal projections rather than neuronal somata in the most superficial cortical layer. These results demonstrate expression of both D1R and D2R in PFC astrocytes, suggesting that PFC astrocytes may respond specifically to DA, as has been demonstrated in other brain regions (Chai et al., 2017; Corkrum et al., 2020; Cui et al., 2016; Fischer et al., 2020; Jennings et al., 2017; Xin et al., 2019).

**Figure 2:**
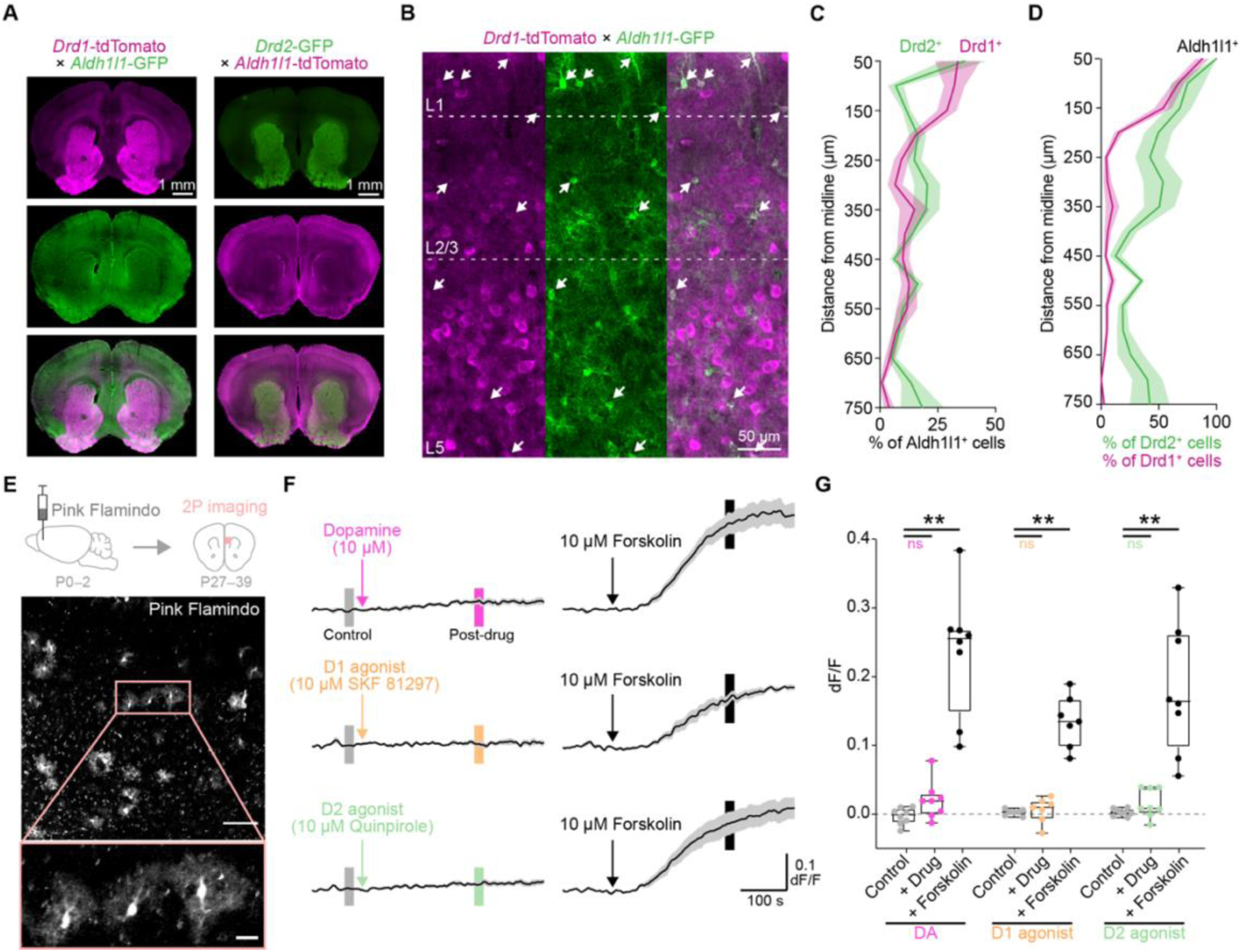
D1 and D2 receptors are expressed by PFC astrocytes but do not recruit G_s_/G_i_ pathways. **(A)** Transgenic crosses to identify co-expression of D1 (left column, top) or D2 receptors (right column, top) with the astrocytic marker Aldh1l1 (middle row). Whole-brain coronal sections ∼1.6 mm anterior to Bregma. **(B)** Example of marker colocalization in PFC of a *Drd1*-tdTomato x *Aldh1l1*-GFP mouse. Arrowheads indicate astrocytes co-expressing D1 (magenta) and Aldh1l1 (green). Boundaries between cortical layers indicated by dashed lines for reference. **(C–D)** Percentage of (C) Aldh1l1^+^ astrocytes expressing D1 (Drd1^+^, magenta) and D2 (Drd2^+^, green), and of (D) Drd1^+^ (magenta) and Drd2^+^ (green) cells that co-express Aldh1l1 in PFC. Data shown as mean ± sem; n = 3 sections/mouse, 3 (D1) and 2 (D2) mice. **(E)** Experimental schematic for 2P imaging of astrocytic cAMP in acute PFC slices. Micrographs show Pink Flamindo expression in entire imaged field-of-view (top), and 3 zoomed-in cells with clear astrocyte morphology (bottom). Scale bars = 100 μm (top) and 20 μm (bottom). **(F)** Dopamine receptor agonists (colors) do not mobilize whole-cell cAMP signaling in PFC astrocytes. Adenylate cyclase activator Forskolin (black) was used in the same slices to confirm Pink Flamindo activity. Neuronal action potentials and neuron-to-astrocyte communication were prevented using TTX and a drug cocktail (see Methods). Vertical boxes on traces indicate 20-s windows used for quantification in (G). Traces shown as slice averages ± sem of whole-cell changes in Pink Flamindo intensity (dF/F); n = 110–180 cells, 7–8 slices, 7–8 mice. **(G)** Quantification of (F) at time-points indicated by small vertical boxes, shown as box plots indicating mean and 10^th^‒90^th^ percentile, and error bars indicating minima and maxima. Slice mean ± sem (Control, +Drug, +Forskolin): -0.003±0.004, 0.02±0.01, 0.24±0.03 (DA); 0.003±0.002, 0.006±0.007, 0.13±0.01 (D1); 0.003±0.002, 0.016±0.007, 0.18±0.03 (D2). Friedman test after Levene test; n.s., *p* > 0.05, **, *p* < 0.01; not shown on graph are comparison between Drug and Forskolin (*p* < 0.05 for all agonists), and comparisons within conditions (controls, Drugs, or Forskolins; one-way Anova or Kruskal-Wallis after Levene test, all *p* > 0.05); n = 110–180 cells, 7–8 slices, 7–8 mice.

### Direct DA receptor stimulation does not recruit the canonical cAMP intracellular signaling pathway

D1R and D2R are canonically coupled to G_s_- and G_i_-GPCR proteins, respectively. To test whether these receptors in PFC astrocytes lead to changes in intracellular cAMP when activated, we expressed the fluorescent cAMP reporter Pink Flamindo (Harada et al., 2017) in PFC astrocytes and performed *ex vivo* acute slice experiments from young adult mice (Fig 2E), while pharmacologically targeting DA receptors (Fig 2F–G). We blocked possible contributions from neighboring D1R^+^ and D2R^+^ neurons by inhibiting action potentials with TTX and preventing neuron-to-astrocyte communication with a multi-drug cocktail (see Methods). We first bath-applied 10 μM DA (Fig 2F left, top; doses in these experiments were chosen to reflect physiological levels of DA reported in the literature), and did not observe significant changes in average Pink Flamindo fluorescence (Fig 2G). However, because D1R and D2R have opposing effects on adenylate cyclase (AC), DA could in principle both stimulate and inhibit cAMP production. Thus, we searched at the single-cell level for both increases and decreases in Pink Flamindo fluorescence, and still did not observe changes after addition of DA (Fig S2B).

To expand on this unexpected negative result and distinguish between possible contributions of G_s_ and G_i_ signaling, we next directly activated either D1R or D2R with receptor subtype-specific agonists (D1: SKF81297, 10 μM; D2: Quinpirole, 10 μM) and imaged cAMP (Fig 2F left, middle–bottom). Again, we found no change from baseline in average cAMP levels for any of the drugs tested (Fig 2G). (Quinpirole is a full agonist at all D2-like dopamine receptors (D2, D3 and D4). However, because all D2-like receptors are coupled to G_s_ proteins—thus canonically linked to increases in cAMP levels—we assumed that this widely used D2 agonist would cause similar changes in cAMP regardless of the receptor subtype involved. Because no response to Quinpirole was detected, we did not explore this further.) To confirm that Pink Flamindo could detect changes in cAMP in our experiments, we followed each drug-treatment experiment with bath-application of the AC agonist Forskolin (10 μM, Fig 2F right). Forskolin application led to a consistent increase in Pink Flamindo fluorescence relative to both baseline and drug treatment (Fig 2G and Fig S2B), which was comparable to Forskolin stimulation in naïve slices (in the absence of TTX and drug cocktail; Fig S2C). We confirmed that these results were not due to slice-to-slice variability (within-treatment comparisons, not significant; Fig 2G) or cell-to-cell differences in Pink Flamindo expression (Fig S2D), indicating that neither DA nor DR subtype-specific agonists induce detectable changes downstream of G_s_ or G_i_ effector proteins in PFC astrocytes.

### DA activates PFC astrocyte Ca^2+^ signals via cell-surface adrenergic receptors

While DRs on PFC may not recruit the cAMP pathway in our experimental conditions, previous research demonstrates 1) that DA can mobilize intracellular Ca^2+^ in astrocytes in hippocampus (Chai et al., 2017; Jennings et al., 2017), olfactory bulb (Fischer et al., 2020) and subcortical nuclei (Chai et al., 2017; Corkrum et al., 2020; Cui et al., 2016; Xin et al., 2019) and 2) DRs in neurons can activate non-canonical GPCR signaling pathways which result in Ca^2+^ mobilization (Lee et al., 2004; Medvedev et al., 2013; Sahu et al., 2009; Undie and Friedman, 1990). To test whether DA mobilizes intracellular Ca^2+^ in PFC astrocytes—and may be mediating the *in vivo* Ca^2+^ activity we described (Fig 1)—we expressed GCaMP6f in PFC astrocytes using viral vectors (Fig 3A–B), and carried out bath-application imaging experiments in acute slices, blocking neuronal activity with TTX and the drug cocktail as above. We found that bath-application of DA caused a large increase in astrocyte Ca^2+^ event frequency compared to baseline (Movie 2; Fig 3C–D, and Fig 3E pink). In contrast, application of D1 and D2 receptor agonists (SKF38393 and Quinpirole, 10 μM) had no discernible effect on Ca^2+^ activity (Fig 3E, yellow; Fig S3A top left). To test whether DRs are engaged in mediating the DA-dependent increase in intracellular Ca^2+^, we next bath-applied DA in the presence of the DR antagonists SCH23390 and Sulpiride, and observed a partial inhibition of astrocyte Ca^2+^ dynamics (Fig S3A, top right), although no significant decrease in event rate compared to application of DA alone (Fig 3D, blue).

**Figure 3:**
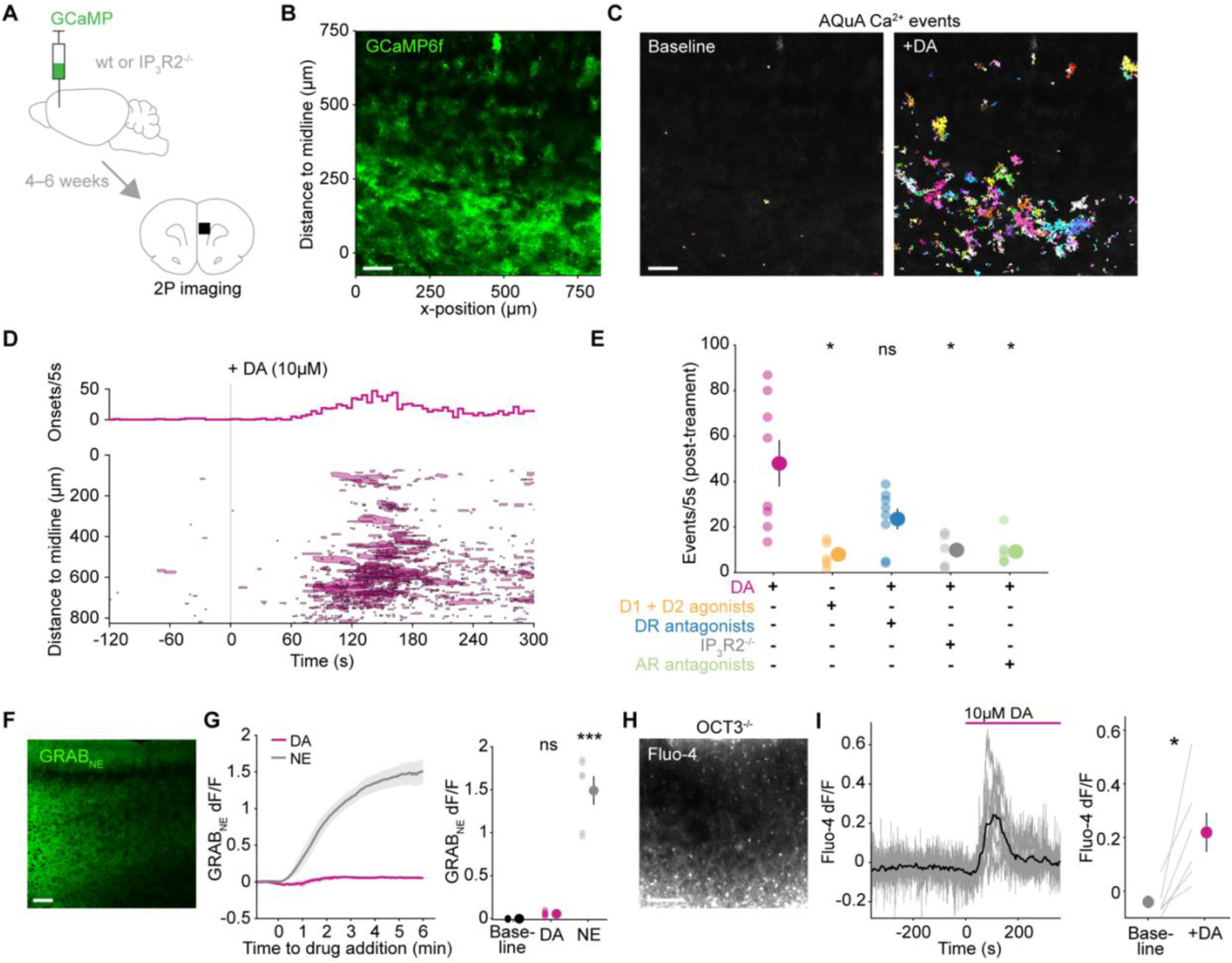
DA mobilizes astrocytic Ca^2+^ in PFC slices via cell-surface adrenergic receptors. **(A)** Experimental strategy for 2P imaging of astrocytic Ca^2+^ in acute PFC slices. **(B)** Representative micrograph of *GfaABC_1_D*-GCaMP6f expression in imaged area of PFC. Note y-axis measurements of distance to slice midline, which are used for spatial plots in (D). Scale bar = 100 μm throughout figure **(C)** All AQuA-detected Ca^2+^ events 0–60 s before (left) and 90–150 s after bath-application of DA (right) from same slice as in (B). Colors represent individual events. **(D)** Time course of all Ca^2+^ events detected over the entire recording of slice in (B–C) and event onset rate (top) relative to 10 μM DA (t = 0). Shaded areas represent approximate event size and mean event y-position over time. **(E)** Astrocytic Ca^2+^-event rate (count/5 s) in PFC slices after treatment with indicated drugs. Treatment with both D1 and D2 agonists SKF38393 and Quinpirole (yellow) did not have the same effect of inducing increased Ca^2+^ events as DA (magenta). Blocking both D1 and D2 receptors with DRs antagonists SCH23390 and Sulpiride during DA application (blue) failed to occlude astrocyte activation by DA, whereas the effect of DA alone was significantly reduced in IP_3_R2^-/-^ mice (grey). Likewise, non-selective α- and β-AR antagonists Phentolamine and Propranolol (green) significantly reduced the astrocytic Ca^2+^ response to DA. Data shown as slices (transparent dots) and corresponding mean ± sem (solid dot and error bar): 48.0±10.2 (DA); 8.0±2.5 (D1/D2 ago.); 23.6±4.6 (DR antag.); 9.9±3.3 (IP_3_R2^-/-^); 9.3±2.5 (AR antag.). Kruskal-Wallis test after Levene test; *, *p* < 0.05 compared to DA condition; all other comparisons between conditions (not shown), *p* > 0.79; n = 5–8 slices, 4–8 mice. **(F)** Example of tissue-wide expression of GRAB_NE_ for 2P imaging in an acute PFC slice. **(G)** DA is not metabolized to NE in PFC slices, as indicated by GRAB_NE_ dynamics. Left: trace means ± sem relative to either DA or NE addition at t = 0. Right: slices (dots) and mean ± sem (dot with error bar) of 20-s GRAB_NE_ dF/F averages at baseline (black), or 6 min after 10 μM DA (magenta) and 10 μM NE (grey): -0.002±0.003 (Baseline); 0.055±0.012 (DA); 1.490±0.165 (NE). Kruskal-Wallis test after Levene test; ***, *p* < 0.001 relative to baseline; n = 6 slices, 4 mice. **(H)** Example acute slice from OCT3^-/-^ mice with deficient DA uptake, loaded with the Ca^2+^ indicator Fluo-4. **(I)** Somatic Ca^2+^ signals in response to bath-applied DA are present in PFC astrocytes in the OCT3^-/-^ background. Left: mean trace (black) and slice average traces of active cells (grey) relative to addition of 10 μM DA at t = 0. Right: slice averages (lines) and corresponding mean ± sem (dots and error bars) of Fluo-4 dF/F extracted from traces on left at either 100-s before (Basal) or after DA (+DA): - 0.04±0.02 (Baseline); 0.22±0.08 (+DA). Paired t-test after Anderson-Darling test; *, *p* < 0.05; n = 138 active cells, 6 slices, 3 mice.

Since the effect of DA on PFC astrocyte Ca^2+^ is minimally inhibited by antagonists of DRs, we next tested whether the robust Ca^2+^ response to DA is mediated by GPCRs at all. To do so, we carried out DA-application and GCaMP imaging experiments in slices obtained from mice genetically lacking IP_3_R2 (Li et al., 2005), the main intracellular receptor downstream of GPCRs in astrocytes that mediates intracellular Ca^2+^ release (Petravicz et al., 2008; Zhang et al., 2014). In these slices, we observed a significant inhibition of Ca^2+^ mobilization by DA (Fig 3D, gray and Fig S3A, bottom left), suggesting that PFC astrocytes do indeed rely on GPCR signaling to mediate the Ca^2+^ response to DA. Because DA has been shown to act on adrenergic receptors (ARs) in neurons (Alachkar et al., 2010; Cilz et al., 2014; Cornil et al., 2002; Guiard et al., 2008; Marek and Aghajanian, 1999; Özkan et al., 2017), we next carried out DA-application in the presence of broad-spectrum adrenergic receptor antagonists (α1/α2: Phentolamine; β: Propranolol; 10 μM). In contrast to the effect of DRs antagonists, blocking ARs completely abolished the DA-mediated increase in Ca^2+^ activity (Fig 3C, green and Fig S3A, bottom right).

Because the astrocyte recruitment by bathed DA had an onset on the order of minutes and was sensitive to ARs inhibitors, we hypothesized that DA could be transformed to NE, a one-step enzymatic product of DA (Kirshner, 1957). To confirm that PFC astrocytes were indeed responding to DA and not NE, we imaged acute PFC slices in which the fluorescent sensor GRAB_NE_ was expressed throughout the tissue (Fig 3F), and bath-applied either DA or NE. In these experiments, DA did not induce a significant change in GRAB_NE_ fluorescence compared to the baseline period, in contrast to a large response to NE (Fig 3G), suggesting that the observed response to DA mediated by ARs (Fig 3C) is not linked to conversion of DA to NE *in situ*. Lastly, we wanted to test whether DA induced PFC astrocyte Ca^2+^ via GPCR signaling from the plasma membrane, or via intracellular compartments, since GPCR signaling can occur or be amplified on internal organelles (Calebiro et al., 2009; Irannejad et al., 2013; Kotowski et al., 2011) and dopaminergic antagonists can display low membrane-permeability (Dos Santos Pereira et al., 2014). To do this, we imaged PFC astrocyte Ca^2+^ (using the indicator Fluo-4) in organic cation transporter 3 knockout mice (OCT3^-/-^, Fig 3H), in which the intracellular transport of monoamines including DA is blocked (Amphoux et al., 2006; Cui et al., 2009; Duan and Wang, 2010; Zwart et al., 2001). Bath-application of DA in slices from these animals (Fig 3I left) led to a robust increase in Fluo-4 fluorescence compared to baseline (Fig 3I right), suggesting that DA acts on cell-surface GPCRs in PFC astrocytes.

### Physiological concentrations of DA evoke fast, transient Ca^2+^ signals in PFC astrocytes

The previous experiments demonstrated that PFC astrocytes respond to continuous application of DA with increases in intracellular Ca^2+^, although with a slow onset. We next explored whether astrocytes can also be engaged by acute stimuli that better reflect physiological dynamics of DA in the brain. To do so, we turned to one-photon (1P) photoactivation of a caged form of DA (RuBi-DA, Fig 4A–B) (Araya et al., 2013), which allowed us to achieve fast release and to mimic volume transmission (Agnati et al., 1995; Banerjee et al., 2020), the main modality of DA release in PFC. We first validated our light-stimulation protocol by comparing a DA dose-response curve (Fig 4C and Fig S4A) to photoactivation of RuBi-DA (Fig 4D and Fig S4B left) in PFC slices expressing the fluorescent DA sensor dLight (Patriarchi et al., 2018). We used these methods to estimate the concentration of DA across the imaging field, and found that our photoactivation protocol released on average ∼2 µM DA (Fig 4E), which matches physiological levels of DA detected by voltammetry in PFC *in vivo* (Garris and Wightman, 1994) and DA concentration estimates near release sites in other brain areas (Courtney and Ford, 2014; Patriarchi et al., 2018).

**Figure 4:**
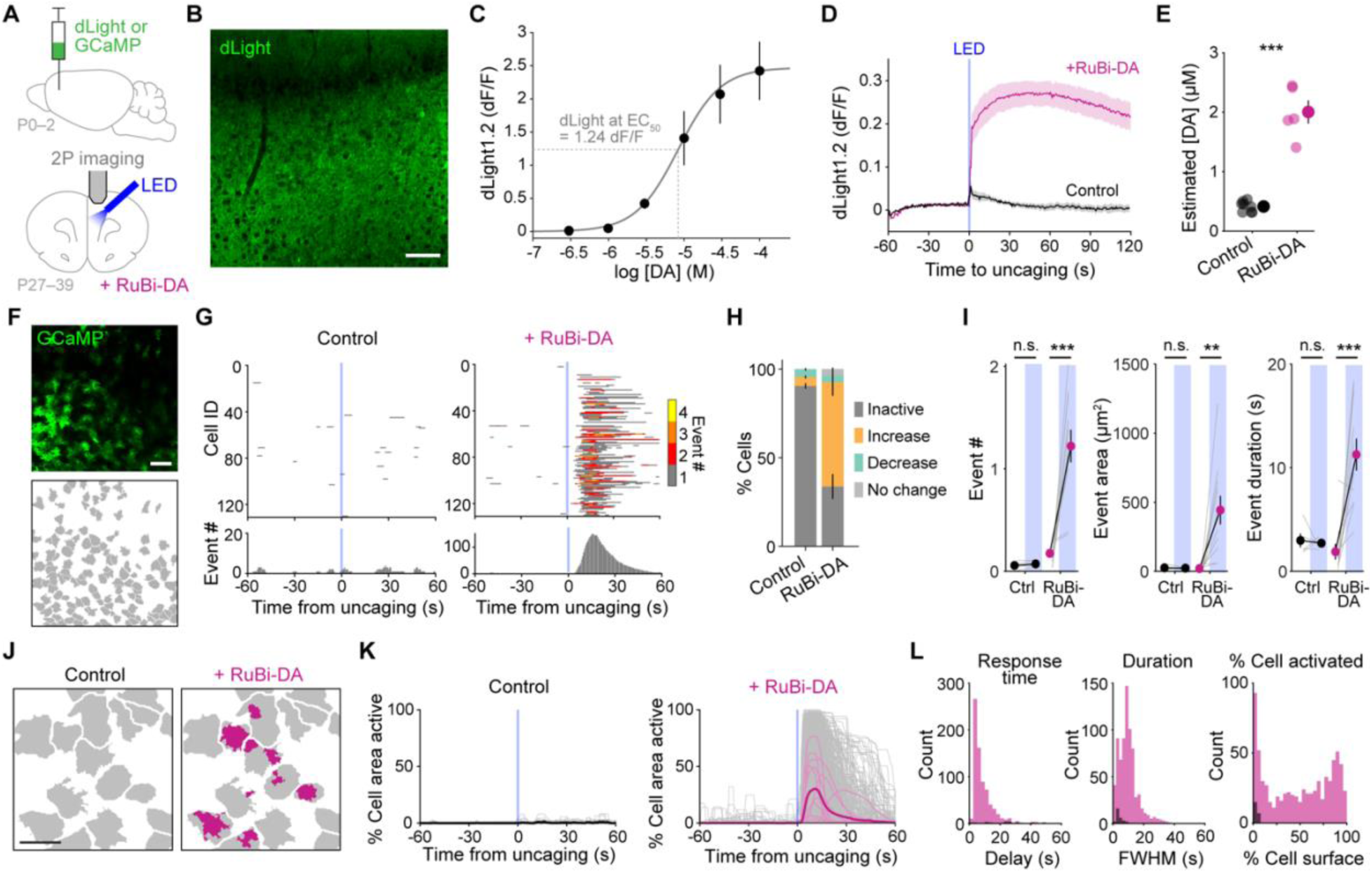
Photo-uncaging in PFC releases physiological concentrations of DA and activates astrocyte territories within seconds. **(A)** Experimental strategy for fast release of DA in PFC slices using RuBi-DA uncaging with a blue LED, combined with simultaneous 2P imaging of DA (dLight1.2) or astrocytic Ca^2+^ (GCaMP6f). **(B)** Representative dLight expression with 2P imaging in an acute PFC slice. Scale bar = 100 μm. **(C)** Dose-response curve of dLight in PFC slices challenged with increasing concentrations of DA, and Hill equation fit function (solid line). Dotted lines indicate DA concentration at dLight half-maximum. Data shown as slice means ± sem (dots and error bars); n = 4 slices, 2 mice. **(D)** dLight fluorescence (dF/F) increases after LED stimulation (3x 100-ms pulses, blue line) in presence of the caged compound RuBi-DA (magenta) but not in th e control without RuBi-DA in the bath (grey). Trace mean ± sem; n = 6–7 slices, 3 mice. **(E)** Estimated DA concentration after RuBi-DA uncaging was 2 μM, extrapolated from fit function in (C) using data from (D) obtained as 30-s dF/F means after LED stimulation. Data shown as all slices (transparent dots) and corresponding mean ± sem (solid dot and error bar): 0.41±0.03 (Control), 2.0±0.2 (RuBi-DA) μM. Two-sample t-test, ***, *p* < 10^-6^; n = 6–7 slices, 3 mice. **(F)** Representative PFC slice expressing astrocytic GCaMP during 2P Ca^2+^ imaging (top) with corresponding cell maps drawn as boundaries around territories of individual astrocytes (bottom). Scale bar = 100 μm. **(G)** Raster-plots of AQuA-detected Ca^2+^ events (top) show the time course of events within individual cells from slice in (F), plotted relative to LED stimulation (blue lines, t = 0) before (left, control) and after bathing on RuBi-DA (right). Colors indicate co-occurring event number in each cell. Cumulative event counts across cells shown in bottom graphs. **(H)** Astrocyte Ca^2+^ activity increases for the majority of cells in the 60 s following RuBi-DA uncaging (70%), while cells are largely inactive (no Ca^2+^ events throughout the recording; 91%) in the control condition. Percentage of cells decreasing or maintaining (no change) their activity after uncaging is similar between conditions, indicating that DA does not induce a decrease in activity in a subset of astrocytes. Data shown as mean ± sem; n = 540–1118 cells, 5–11 slices, 5–8 mice. % Cells (Control, RuBi-DA): 91±2, 30±7 (Inactive); 4±1, 62±8 (Increase); 4±1, 4±1 (Decrease); 0±0, 4±1 (No change). **(I)** Ca^2+^ event features (number, area, duration) in active cells in (H) increase significantly in the 60 s after uncaging light (shaded blue boxes) with RuBi-DA (magenta) but not without (control, black). Slice averages of active cells (grey lines) and mean ± sem (dots and error bars). Event # (pre-, post-uncaging): 0.06±0.02, 0.07±0.01 (Control); 0.17±0.03, 1.22±0.16 (RuBi-DA). Event area (µm^2^): 25±7, 23±3 (Control); 20±5, 443±104 (RuBi-DA). Event duration (s): 3.0±0.7, 2.7±0.2 (Control); 1.9±0.8, 11.3±1.5 (RuBi-DA). Paired t-test comparing pre- to post-uncaging; **, *p* < 0.01; ***, *p* < 0.001. Control: n = 47/540 active/total cells, 5 slices and mice. RuBi-DA: n = 784/1118 cells, 11 slices, 8 mice. **(J)** Example of Ca^2+^ activation within individual cells 5 s after the uncaging pulse, either in control (left) or after addition of RuBi-DA (right). Maps are zoomed in from maps in (F). Grey = cell areas; magenta = active pixels. Scale bar = 50 μm. **(K)** Time course of % cell area active relative to uncaging (blue lines) in absence (control) and presence of RuBi-DA. Data shown as cells, slices, and overall mean (grey, thin and thick colored lines, respectively). Control: n = 47/540 cells, 5 slices and mice. RuBi-DA: n = 784/1118 cells, 11 slices, 8 mice. **(L)** Astrocytic response to DA release (magenta) occurs within seconds of LED stimulation (left, delay), lasts <20 s (middle, peak full-width half-maximum), and recruits a wide range of areas within individual astrocytes (right, % cell surface). In controls (black), few cells were active after uncaging, with short activity (< 9 s) covering a small percentage of cell area. Control: n = 22 cells, 5 slices, 5 mice. RuBi-DA: n = 720 cells, 11 slices, 8 mice.

Next, to understand how single PFC astrocytes respond to temporally controlled DA release, we uncaged RuBi-DA in slices with GCaMP-expressing astrocytes (Fig 4F top, Movie 3) while blocking neurons with TTX and drug cocktail as above. For analysis, we drew borders around each cell (Fig 4F, bottom) and detected Ca^2+^ events using AQuA within these single-cell maps, which allowed us to map Ca^2+^ event features within cells (Fig 4G–I) and monitor the amount of cell recruited over time (Fig 4J–L) in response to temporally precise DA release. In control conditions (no RuBi-DA; Fig 4G–H, left), the majority of astrocytes (91%) were inactive throughout the trial and similar numbers of cells increased or decreased their Ca^2+^ event activity around the LED pulse (4%). In contrast, in presence of RuBi-DA (Fig 4G–H, right), the majority of astrocytes across all cortical layers responded to uncaging with an increase in activity (62%) rather than a decrease (4%). We next explored the Ca^2+^ event features induced by uncaging in individual astrocytes, and found that events were more abundant, larger, and lasted longer following LED stimulation in RuBi-DA, but not in the controls (Fig 4I). We confirmed that these results were not affected by the pharmacological cocktail used since all features of Ca^2+^ events were unchanged compared to naïve slices (Fig S4C–D).

When looking at the overall Ca^2+^ mobilization within individual astrocytes (Fig 4K), we observed that Ca^2+^ activity was induced with a short response time (8.6 s, Fig 4K, L, left) and short duration (9.9 s, Fig 4K, middle) in most cells, whereas the amount of cell recruited by fast DA release varied considerably between cells (49%, Fig 4K, right). We confirmed that the results we obtained were not affected by our single-cell delineation method, as we found no correlation between cell size and amount of cell surface recruited by DA (Fig S4E). These data demonstrate that astrocytes respond acutely to physiological levels of extracellular DA with fast and transient Ca^2+^ dynamics that cover different astrocyte territories across cells.

### PFC astrocytes require α1-AR signaling to respond to DA

Our bath application experiments suggested that ARs—but not DRs—mediate signaling downstream of DA in PFC astrocytes (Fig 3E). To explore the contribution of DRs and ARs to fast dopaminergic stimulation of astrocytes, we photoreleased DA as described above on PFC slices treated with different subtype-specific inhibitors of DA and NE receptors (Fig 5A, Movie 4). We monitored population-level astrocyte Ca^2+^ while blocking neuronal contributions, and similar to our previous experiments (Fig 4), we observed a significant increase in astrocyte Ca^2+^ following uncaging of RuBi-DA alone (Fig 5B–C, pink; control). Antagonizing D1 or D2 receptors did not occlude the response to DA release (Fig 5B–C), in accordance with our bath-application data (Fig 3E) and further supporting the idea that DRs are not involved in the recruitment of PFC astrocytes by DA. Next, we tested the contribution of all AR subtypes (α1, α2 and β) to DA-mediated astrocyte Ca^2+^ activity, and found that only inhibition of α1-AR prevented Ca^2+^ mobilization after RuBi-DA uncaging (Fig 5B–C). We also measured astrocyte activity using different metrics (raw Ca^2+^ events dF/F [Fig S5A] and percent of field-of-view recruited [Fig S5B]) and found no change from the above results, further supporting the observation that only α1-ARs are necessary for the activation of PFC astrocytes by DA. Overall, these *ex vivo* data suggest that fast, volume transmission-like release of DA at physiological concentrations recruits PFC astrocytes via α1-ARs.

**Figure 5:**
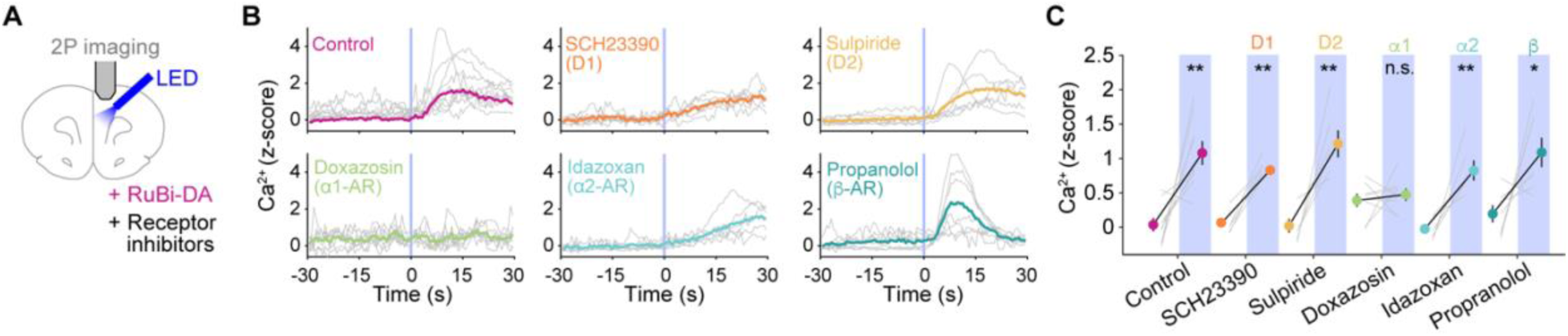
Fast astrocyte responses to DA in PFC slices occur via α1-ARs. **(A)** Experimental schematic for RuBi-DA uncaging and simultaneous 2P imaging of astrocyte Ca^2+^ in PFC slices bathed with different receptor antagonists. **(B)** Astrocyte Ca^2+^ in PFC slices increases shortly after RuBi-DA uncaging (control), an effect blocked by α1-AR antagonist Doxazosin (10 μM), but not by D1 (SCH23390, 10 μM), D2 (Sulpiride, 0.5 μM), α2-AR (Idazoxan, 10 μM) or β-AR (Propranolol, 10 μM) antagonists. Data relative to uncaging (blue lines, t = 0) as slice average traces (grey lines) of AQuA-detected, z-scored Ca^2+^ events in GCaMP6f-expressing astrocytes, with overall mean as colored traces. **(C)** Quantification of (B), shown as 30-s mean of slice Ca^2+^ immediately before (white) or after RuBi-DA uncaging (shaded blue boxes) in presence of receptor inhibitors as indicated by labels. Data shown as slices (grey lines) and corresponding mean ± sem (black lines, solid dots, and error bars): 0.03±0.10, 1.08±0.17 (control); 0.07±0.07, 0.83±0.06 (D1); 0.02±0.10, 1.21±0.20 (D2); 0.39±0.10, 0.47±0.09 (α1); -0.03±0.06, 0.82±0.15 (α2); 0.19±0.12, 1.09±0.21 (β). Paired t-test after Anderson-Darling test to compare pre- to post-uncaging values; *, *p* < 0.05, **, *p* < 0.01; *p* = 0.004 (control), 0.0006 (D1), 0.008 (D2), 0.624 (α1), 0.008 (α2), 0.036 (β); n = 6–9 slices, 5–9 mice. Pre-uncaging values in treatments versus control were not statistically different (Kruskal-Wallis test with Dunn’s correction: adjusted p-value > 0.19 for all comparisons).

### DA evokes Ca^2+^ signals in PFC astrocytes via α1-ARs *in vivo*

Our previous results indicate that DA induces fast, transient Ca^2+^ signals in prefrontal astrocytes in acute slices. To test whether dopaminergic inputs to the PFC induce astrocyte activity *in vivo*, we carried out dual-color fiber photometry recordings using viral expression of the red-shifted Ca^2+^ sensor jR-GECO1b and the DA sensor dLight to monitor both extracellular DA and astrocyte Ca^2+^ dynamics (Fig 6A and Fig S6E). Because aversive stimuli such as foot shock (Thierry et al., 1976), tail shock (Abercrombie et al., 1989), and tail pinch (Vander Weele et al., 2018) are known to activate the mesocortical DA system, we used an aversive tail-lift stimulus (Hurst and West, 2010) to increase DA levels in PFC. Using this experimental paradigm and monitoring extracellular DA concentration using dLight, we found that this was indeed the case (Fig 6A, green). jR-GECO1b recordings also showed that there were large astrocyte Ca^2+^ transients during the tail-lift procedure (Fig 6A, pink). We aligned the transients from these two recording channels, and saw that the jR-GECO signal closely followed dLight (Fig 6B–C and Fig S6A). Cross-correlation of the two signals indicated that dLight precedes the jR-GECO signal by 1.4 s (Fig 6D), suggesting that extracellular DA in PFC is a major contributor to the astrocyte Ca^2+^ recruitment we observe during this aversive stimulus.

**Figure 6:**
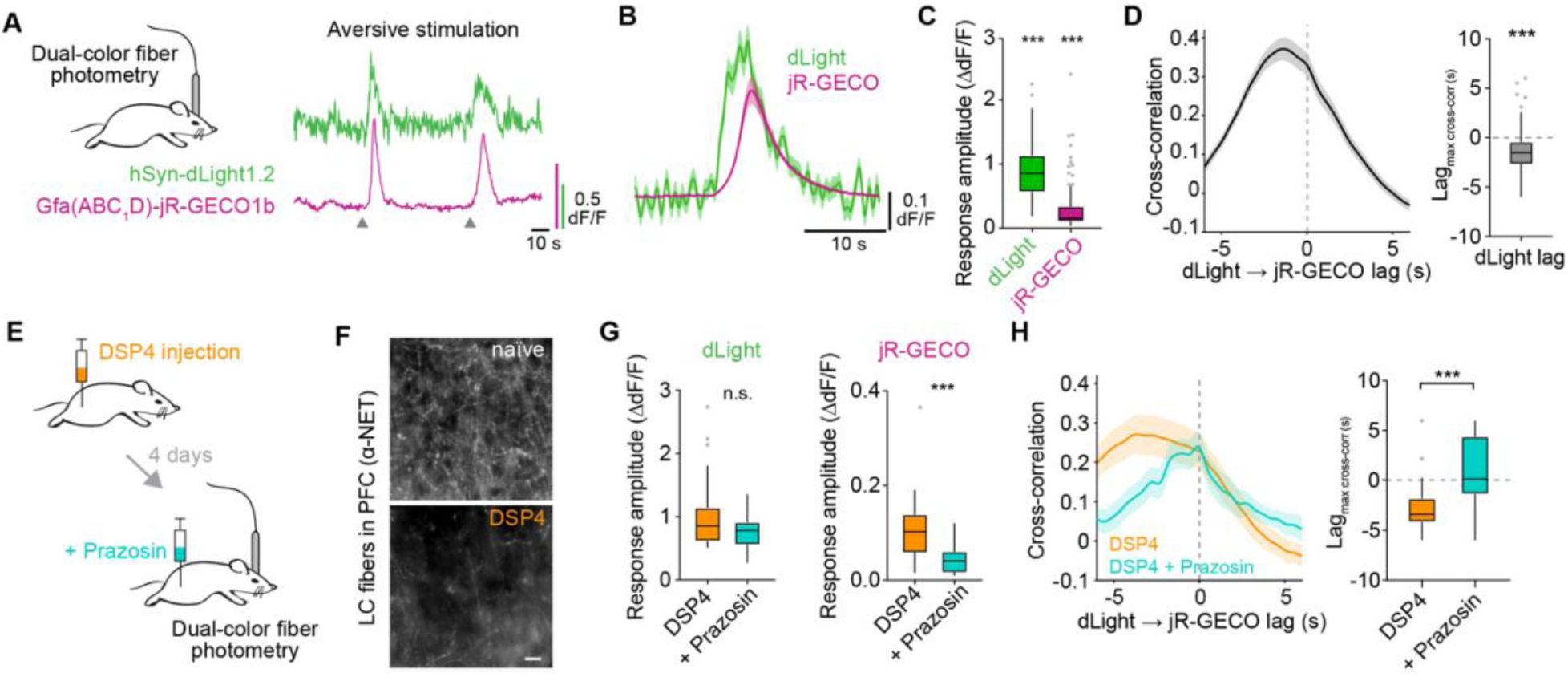
Astrocyte Ca^2+^ follows DA release *in vivo* via α1-ARs. **(A)** Left: experimental setup for dual-color fiber photometry in PFC of behaving mice for recording DA (*hSyn*-dLight1.2, green) and astrocyte Ca^2+^ (*GfaABC_1_D*-jR-GECO1b, magenta) dynamics. Right: example traces during aversive stimulation by tail lift (gray triangles). **(B)** Average photometry traces for DA and astrocyte Ca^2+^ in PFC *in vivo*, aligned to the onset of astrocyte Ca^2+^ transients. Data shown as mean ± sem; n = 96 transients, 9 mice. **(C)** Response amplitude for DA (green) and astrocyte Ca^2+^ (magenta) in aversive stimulation experiments deviate significantly from baseline values (dLight: 0.89±0.04 dF/F; jR-GECO: 0.31±0.04 dF/F). Data shown as Tukey boxplots, calculated as maximum dF/F relative to 20-s mean before the jR-GECO peak. One-sample t-test or sign test with hypothesized mean 0, after Anderson-Darling test to show difference from 0; ***, *p* < 0.001; n = 96 transients, 9 mice. **(D)** Cross-correlation of dLight and jR-GECO traces (left) indicates that DA signals in PFC *in vivo* precede astrocyte Ca^2+^ transients by 1.4±0.2 s (right). Data shown as mean ± sem and Tukey boxplots; one-sample sign test with hypothesized mean 0, ***, *p* < 0.001; n = 96 transients, 9 mice. **(E)** Schematic of dual-color fiber photometry in PFC of mice treated with the LC-toxin DSP4 (50 mg/kg, i.p., 2 injections, 2 days apart), before and after administration of the α1-AR antagonist Prazosin (5 mg/kg, i.p.). **(F)** LC inputs to PFC revealed by NET immunostaining (top, naïve) are decreased after DSP4 treatment (bottom). Scale bar = 50 μm. **(G)** In NE-depleted animals, DA transients (left, dLight) during aversive stimulation in PFC are still present (DSP4, orange; 1.02±0.11 dF/F) and unaffected when α1-ARs are blocked (+ Prazosin, aqua; 0.77±0.05), whereas Ca^2+^ peaks in astrocytes (right, jR-GECO) are significantly reduced by Prazosin treatment (DSP4: 0.11±0.01 dF/F; +Prazosin: 0.04±0.0 dF/F), indicating that astrocyte Ca^2+^ in the PFC rely on α1-ARs even with diminished NE release. Data shown as Tukey boxplots, calculated as maximum dF/F relative to 20-s means before jR-GECO peaks. Wilcoxon rank sum test; ***, *p* = 0.0003; n = 27–29 transients, 4 mice. **(H)** Cross-correlation of dLight and jR-GECO traces (left) in NE-depleted animals (DSP4), and in the same animals after inhibition of α1-ARs (DSP4 + Prazosin) shows that DA signals in PFC precede astrocyte Ca^2+^ with diminished NE (DSP4, -2.64±0.52 s) but not after inhibition of α1-ARs (+Prazosin, 0.73±0.65 s). Data shown as trace mean ± sem and Tukey boxplots; Wilcoxon rank sum test; ***, *p* = 0.0003; n = 27–29 transients, 4 mice.

Because aversive stimulation also releases NE in the PFC (Gresch et al., 1994), we next sought to describe any contribution of NE to this close relationship between DA and astrocyte Ca^2+^ we observed in the PFC *in vivo*. To do this, we carried out similar dual-color fiber photometry experiments as previously, but after injection of DSP4 (Fig 6E), a toxin that specifically ablates locus coeruleus (LC) projection fibers (Bekar et al., 2008; Ding et al., 2013; Fritschy and Grzanna, 1989), the main source of NE release in PFC. We confirmed that this approach reduced LC fibers in the PFC by performing norepinephrine transporter (NET) immunostaining after DSP4 treatment (Fig 6F), and again compared dLight and jR-GECO dynamics. The amplitude of the DA signals in response to our aversion paradigm were unchanged in astrocytes of NE-depleted mice compared to controls (Fig S6B, left), supporting the selectivity of the toxin used in targeting LC fibers. In addition, while we observed a decrease in astrocyte Ca^2+^ amplitude (Fig S6B, right)—in accordance with the documented role of NE in astrocyte Ca^2+^ signaling in other brain regions (Bekar et al., 2008; Ding et al., 2013; Gordon et al., 2005; Paukert et al., 2014)—astrocyte Ca^2+^ transients co-occurring with dLight transients remained evident after NE-depletion (Fig S6A, middle row). These astrocyte Ca^2+^ signals were longer (18 s, Fig S6C), and occurred with longer lag after dLight (2.6 s, Fig S6D) compared to untreated animals (duration 7 s, lag 1.4 s), which may be explained by slower DA uptake in the absence of the NET transporter (Morón et al., 2002; Sesack et al., 1998). Overall, these results indicate that mesocortical DA recruits astrocytes in the PFC when mice are challenged with an aversive stimulation, independent of LC inputs.

Our *ex vivo* experiments indicate that DA effects on PFC astrocytes are mediated by α1-AR (Fig 3, Fig 5). To test whether this functional crosstalk between dopaminergic and noradrenergic systems also occurs *in vivo*, we next compared responses to aversive stimulation in mice treated with the LC toxin DSP4 before and after injection of an α1-AR antagonist (Prazosin, Fig 6G–H). We found that while dLight signals in response to aversion were maintained after Prazosin (Fig 6G, left), astrocyte Ca^2+^ dynamics were significantly reduced (Fig 6G, right) and did not follow DA dynamics (Fig 6H). Together, these data suggest that α1-AR signaling accounts for the bulk of the astrocyte Ca^2+^ response to DA in the PFC *in vivo*.

### DA increases extracellular ATP at PFC astrocytes

Previous work indicates that DA can stimulate ATP release from astrocytes to modulate synaptic transmission in surrounding neurons in *nucleus accumbens* (Corkrum et al., 2020). To answer whether the α1-AR-mediated activation of PFC astrocytes by DA that we observed *in vitro* and *in vivo* also leads to mobilization of extracellular ATP, we performed 2P acute slice experiments on astrocytes expressing the GPCR-based fluorescent ATP sensor (GRAB_ATP_, Fig 7A) (Wu et al., 2021) and detected extracellular ATP events at astrocytes using AQuA (Fig 7B, G) (Wang et al., 2019). We first determined the response dynamics of the sensor in our experimental setup by bathing on exogenous ATP (50 μM, Fig 7B–C), and found that continuous stimulation with ATP led to an increased event rate (Fig 7D), with events whose size matched the territory of an individual astrocyte and could be detected during the entire course of the ATP application (Fig 7E). These data demonstrate that GRAB_ATP_ can reliably detect the presence of ATP over the entire astrocyte surface for prolonged periods of time and is suitable for *ex vivo* imaging of extracellular ATP on PFC astrocytes.

**Figure 7:**
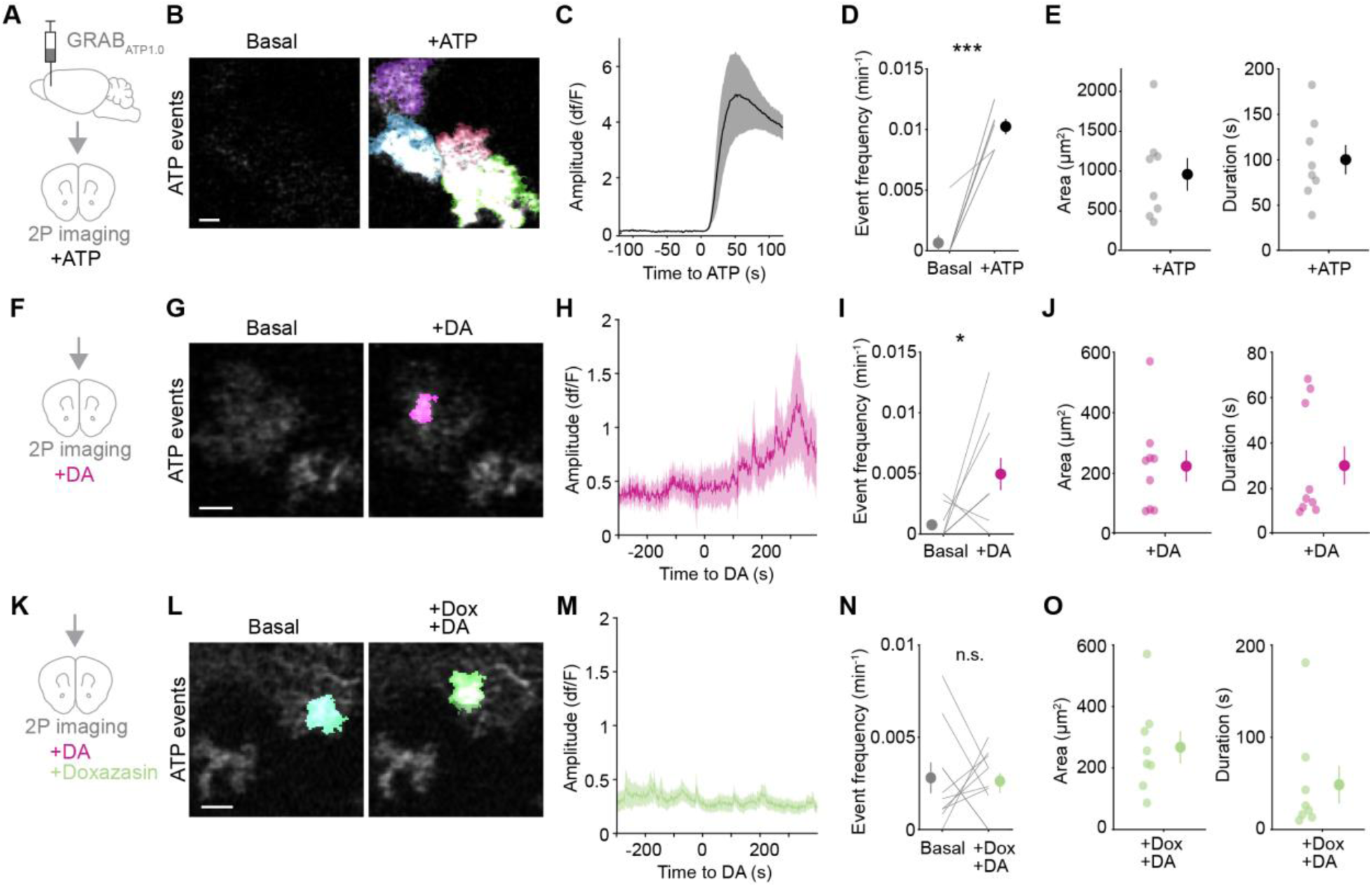
DA mobilizes ATP at discrete locations at PFC astrocytes via α1-ARs. **(A)** Experimental schematic for 2P astrocytic GRAB_ATP_ imaging in acute PFC slices. **(B–C)** Continuous bath-application of ATP (50 μM) induces strong, sustained fluorescent signals in astrocytes, shown as (B) PFC astrocytes expressing GRAB_ATP_ (grayscale) and color overlay of AQuA-detected ATP events before (left, basal) and after ATP (right), and (C) time course of the dF/F amplitude of AQuA-detected ATP events relative to exogenous ATP application (t = 0). Scale bar = 20 μm. Data shown as mean ± sem of slice traces (line and shaded area). n = 52/62 active/total cells, 8 slices, 3 mice. **(D)** GRAB_ATP_ event rate increases following stimulation with ATP. Data shown as slice averages (lines) and mean ± sem (dots and error bars): 0.0007±0.0007 (Basal), 0.010±0.001 (+ATP) min^-1^. Paired t-test after Anderson-Darling test; ***, *p* < 10^-4^; n = 8 slices, 3 mice. **(E)** GRAB_ATP_ events in response to continuous ATP application covered the entire sensor-expressing astrocyte territory (left, 1044±224 μm^2^) and were sustained (right, 100±16 s) as expected. Data shown as slice averages of active cells (transparent dots) and overall mean ± sem (solid dots and error bars); n = 8 slices, 3 mice. **(F–H)** Application of DA (10 μM) onto PFC astrocytes expressing GRAB_ATP_ (F) induces localized ATP events, shown as (G) GRAB_ATP_ micrographs and AQuA overlay, which are delayed (H) as shown by the time course of GRAB_ATP_ event dF/F relative to DA application (t = 0). Scale bar = 20 μm. Data shown as mean ± sem of slice averages (line and shaded area). n = 23/101 active/total cells, 10 slices, 5 mice. **(I)** Rate of ATP events after DA application was higher than baseline. Data shown as slice averages (lines) and mean ± sem (dots and error bars): 0.0007±0.0004 (Basal), 0.005±0.001 (+DA) min^-1^. Paired t-test after Anderson-Darling test; *, *p* = 0.025; n = 10 slices, 5 mice. **(J)** GRAB_ATP_ events in response to DA were smaller than entire astrocyte territories (221±52 μm^2^) and time-restricted (30±8 s). Data shown as slice averages of active cells (transparent dots) and overall mean ± sem (solid dots and error bars); n = 9 slices, 5 mice. **(K–M)** In presence of the α1-AR antagonist Doxazosin (10 μM), DA (K) does not induce ATP events, as shown by (L) GRAB_ATP_ micrographs and AQuA overlay, and (M) time-course of GRAB_ATP_ event dF/F relative to DA application (t = 0, 10 μM). Scale bar = 20 μm. Data shown as mean ± sem of slice averages (line and shaded area). n = 41/160 active/total cells, 10 slices, 5 mice. **(N)** In presence of Doxazosin, DA application does not increase the rate of ATP events. Data shown as slice averages (lines) and mean ± sem (dots and error bars): 0.0028±0.0008 (Basal), 0.0026±0.0005 (+Dox/+DA) min^-1^. Paired t-test after Anderson-Darling test; n.s., *p* = 0.878; n = 10 slices, 5 mice. **(O)** GRAB_ATP_ events in presence of Doxazosin are similar in size (267±53 μm^2^) and duration (48±20 s) compared to those observed in response to DA alone. Data shown as slice averages of active cells (transparent dots) and overall mean ± sem (solid dots and error bars); n = 10 slices, 5 mice.

After validating this approach to detect extracellular ATP at PFC astrocytes, we next asked whether PFC astrocytes stimulated with DA can lead to extracellular ATP mobilization. We repeated ATP imaging experiments while bath-applying DA (10 μM, Fig 7F–H) and blocking neuronal contributions as described above (without PPADS and CGS 15943 to avoid occluding GRAB_ATP_ fluorescence changes, see Methods). We found that DA induces mobilization of extracellular ATP (Fig 7H) and an increase in the frequency of ATP events (Fig 7I). These sparse, DA-induced ATP events lasted ∼30 s and did not encompass the entire astrocyte territory (Fig 7J). These results indicate that ATP is increased at constrained cellular locations at PFC astrocytes in response to DA. We next carried out the same experiments, but added Doxazosin before each recording, to inhibit α1-ARs (Fig 7K). In these experiments, we no longer saw an increase in the frequency of extracellular ATP events after addition of DA (Fig M–N), but the area and duration of the spontaneous events we did see were similar to those observed with addition of DA alone (Fig 7L, O). This result supports the concept that α1-ARs are important for DA signaling that leads to ATP increases. Although we do not rule out the contribution of other, non-astrocytic cell types to this phenomenon, this relationship between DA signaling and ATP may contribute to regulation of synaptic transmission in PFC.

## Discussion

### PFC astrocyte activation relative to behavior

Recent evidence shows that astrocytes play active roles in brain computation and animal behavior (Kol et al., 2020; Martin-Fernandez et al., 2017; Nagai et al., 2019; Poskanzer and Yuste, 2016), including in the PFC (Mederos et al., 2021). In focusing on astrocytic roles in PFC function, we find that PFC astrocytes differ in neurophysiology from astrocytes in sensory cortices (Fig 1). They are activated with different spatiotemporal patterns of intracellular Ca^2+^ (Fig 1), and when animals are exposed to aversive, stressful stimuli (Fig 6), but not in response to locomotion (Fig 1). These results are consistent with PFC neuronal networks being involved in stress processing (Abercrombie et al., 1989; Lammel et al., 2012; Rosenkranz and Grace, 2001; Thierry et al., 1976; Vander Weele et al., 2018), and with studies demonstrating structural, transcriptomic, and metabolic changes in astrocytes following exposure to acute and chronic stress (Abbink et al., 2019; Bender et al., 2020; Murphy-Royal et al., 2020; Simard et al., 2018). Overall, our results are consistent with previous work showing divergent transcriptomic, morphological, and cellular signaling landscapes in astrocytes of different brain areas (Batiuk et al., 2020; Chai et al., 2017; Khakh and Sofroniew, 2015; Xin et al., 2019), and support the hypothesis that astrocytes may serve specific computational or behavioral functions in PFC.

### DA actions on PFC astrocytes: sustained and heterogeneous responses

Sparser input, faster firing (Lammel et al., 2008), and lower uptake (Sesack et al., 1998) cause greater diffusion and prolonged availability of DA in PFC compared to subcortical areas (Abercrombie et al., 1989; Garris and Wightman, 1994), which, combined with tonic and phasic modes of DA release, results in complex effects on neuronal circuit dynamics (Lohani et al., 2019). While the historical focus of DA signaling in PFC has been on neurons, other cell types such as astrocytes are also well positioned to be reached by these sustained, long-range DA signals and thus contribute to regulation of synaptic transmission in PFC. The ability of astrocytes to respond to dopaminergic signaling with Ca^2+^ elevations had been shown in brain areas other than cerebral cortex (Chai et al., 2017; Corkrum et al., 2020; Cui et al., 2016; Fischer et al., 2020; Jennings et al., 2017; Xin et al., 2019). Our study expands this knowledge, demonstrating that astrocytes are capable of sensing DA directly in the PFC, responding with complex Ca^2+^ events to both continuous application (Fig 3) and phasic release (Fig 4) of DA. The different dynamics of PFC astrocyte Ca^2+^ observed in response to these two modalities of DA delivery suggest a possible mechanism by which astrocytes may discern between tonic and phasic DA signals. Importantly, since we blocked action potentials and neuronally released molecules known to bind GPCRs on astrocytes in our *ex vivo* experiments, our data demonstrate that PFC astrocytes respond to DA independently of neuronal activation by DA, suggesting that astrocytes actively contribute to modulation of PFC activity via DA.

Our uncaging data (Fig 4) show that, even in the absence of neuronal responses to DA, the rapid release of physiological concentrations of DA recruits astrocyte responses within seconds, which are sustained for tens of seconds in the majority of cells, but not all. Individual astrocyte responses, rather than population-wide activity, demonstrate that the extent of subcellular locations engaged in Ca^2+^ signaling following DA release within individual cells differs across PFC astrocytes. These observations suggest that astrocytes may be involved in regulation of sustained activity or network dynamics, as well as contribute to the local computation of PFC circuitry in a cell-specific manner, as shown previously in other brain areas (Martín et al., 2015; Martin-Fernandez et al., 2017).

### DA actions on PFC astrocytes: receptors and signaling pathways

Our results demonstrate that DA acting on PFC astrocytes recruits Ca^2+^ (Fig 3–4) rather than cAMP (Fig 2). These data are in contrast with previous neuronal research showing that DA activates the G_s_/G_i_-cAMP pathways canonically ascribed to D1 and D2 receptors (Lee et al., 2021; Muntean et al., 2018; Nomura et al., 2014; Yapo et al., 2017), but in in agreement with previous astrocytic work from other brain areas which have shown that DA activates the G_q_-IP_3_-Ca^2+^ pathway (Chai et al., 2017; Corkrum et al., 2020; Cui et al., 2016; Fischer et al., 2020; Jennings et al., 2017; Xin et al., 2019). Together, these results may suggest differential expression of components of the signaling machinery across brain cell types. However, the possibility supported by our pharmacology data (Fig 3, 5–6) is that the lack of cAMP mobilization is due to DA acting on PFC astrocytes exclusively through α1-AR—the only G_q_-coupled receptor across all DR and AR subtypes—even though PFC astrocytes do express D1 and D2 receptors (Fig 2). Indeed, our data demonstrating that PFC astrocytes respond to DA with fast, transient Ca^2+^ mobilization mediated by α1-ARs differs from previous work on astrocytes in other brain regions, in which DA can induce Ca^2+^ responses via DRs (Corkrum et al., 2020; Fischer et al., 2020; Jennings et al., 2017). However, our data that astrocytes sense DA via cell-surface α1-ARs, and not DRs, is consistent with studies of neuronal activation by DA also report that DRs agonists or antagonists are unable to reproduce or prevent the effects of DA (Cilz et al., 2014; Cornil and Ball, 2008; Cornil et al., 2002; Guiard et al., 2008; Marek and Aghajanian, 1999; Nicola and Malenka, 1997; Özkan et al., 2017). Further, our unexpected finding could help reconcile some apparently contradictory findings in the astrocyte literature whereby different methods (DA versus D1/D2 agonists, or blockade of DA with DR antagonists) for DA activation lead to contrasting results even within brain regions (Corkrum et al., 2020; D’Ascenzo et al., 2007). For instance, the documented dorso-ventral or layer-specific gradients of VTA/LC innervation or DR/AR expression in the hippocampus (Edelmann and Lessmann, 2018) could have influenced the pharmacology of cellular responses observed by Jennings *et al*., whereby lower local expression of DRs in *stratum lacunosum-moleculare* could have allowed AR-mediated DA responses to take over and explain the lack of sensitivity of DA-induced responses to DR antagonists. Similarly, because the transcriptomic, morphological, and cellular signaling landscape of astrocytes can diverge in different cortical layers or brain areas (Batiuk et al., 2020; Chai et al., 2017; Lanjakornsiripan et al., 2018; Xin et al., 2019), region- or subregion-specific patterns of innervation and receptor expression could favor different mechanisms of DA activation—via DRs or ARs—and explain the lack of activation by D1/D2 agonists observed by some studies (Chai et al., 2017; D’Ascenzo et al., 2007), as well as the lack of inhibition by DRs antagonists found by others (Jennings et al., 2017).

### What is the adaptive role of DA/α1-AR promiscuity?

An important implication of our findings is that DA signaling *ex vivo* (Fig 3, 5) and *in vivo* (Fig 6) is subject to receptor promiscuity. This result is supported by previous research reporting that the effects of DA on neurons could not be reproduced using DA-selective agonists (Cilz et al., 2014; Nicola and Malenka, 1997; Özkan et al., 2017), or could be prevented by α-AR, but not DR, antagonists (Cilz et al., 2014; Cornil et al., 2002; Guiard et al., 2008; Marek and Aghajanian, 1999; Özkan et al., 2017). Further, many levels of interactions between the dopaminergic and noradrenergic systems have been documented in the PFC: DA and NE are co-released by LC fibers (Devoto et al., 2005), DA uptake is accomplished mainly by NET (Morón et al., 2002) due to low DAT expression in PFC (Sesack et al., 1998), and sub- or supra-threshold stimulation of both systems leads to detrimental outcomes on PFC performance (Arnsten et al., 2012). Together, this evidence supports the idea that DA may interact with the noradrenergic system at the receptor and signal transduction level on PFC astrocytes. However, despite likely acting through the same astrocytic receptors, DA and NE show markedly different Ca^2+^ mobilization signatures: DA evokes events that are small in amplitude and duration, but occur at high frequency and are prolonged over time (Fig 3), while previous work shows that NE causes strong events of big amplitude and short duration (Pankratov and Lalo, 2015). Thus, the DA-AR crosstalk described in the present study does not implicate losing information about where a given extracellular signal originated from, as astrocytes may still be able to implement different computations and activate different effects downstream of specific inputs, likely through different combinations of receptors recruited and their stoichiometry and position relative to signal transducers and effectors.

Although VTA-DA and LC-NE traditionally play dissociable roles in the brain, evidence of a functional relevance of DA/NE co-release by LC fibers to memory consolidation in the hippocampus (Kempadoo et al., 2016; Smith and Greene, 2012; Takeuchi et al., 2016) could suggest that DA/NE co-release in the PFC (Devoto et al., 2005) might also be relevant to specific prefrontal cognitive processes. However, our data shows that substantial PFC DA dynamics are maintained in the PFC of DSP4-treated mice that lack LC innervation (Fig 6), in accordance with previous work *in vivo* (Berger et al., 1974; Vander Weele et al., 2018). Nevertheless, the specific functional effects of DA/NE co-release in the PFC remain to be explored, and recent advances in extracellular neurotransmitter imaging probes (Feng et al., 2019; Muller et al., 2014; Patriarchi et al., 2018, 2020; Sun et al., 2018) will help understand how astrocytes, as well as neurons, are regulated differentially by these two neuromodulators. For instance, is the release of VTA-DA and LC-DA in PFC driven by similar or divergent behavioral stimuli, leading then to mutual reinforcement or rather reciprocal modulation of segregate inputs? Does co-release of DA/NE from LC fibers in the PFC mediate specific aspects of PFC executive functions? On the other hand, given DA availability in PFC is prolonged compared to that of NE (Devoto et al., 2005) thus leading to greater spatial diffusion, do LC-DA and LC-NE reach similar or different cellular targets, and with what temporal mismatch? In both scenarios, astrocytes are well positioned to detect both temporally segregated or coincident DA/NE signals, as they express receptors for both neurotransmitters whose stimulation generates distinctive cellular responses. Further studies are needed to address the extent of the crosstalk between the DA and NE system, both within astrocytes and across brain cell types, as well as its implications on brain function and animal behavior.

How this receptor/neuromodulator promiscuity originates at the receptor level is another exciting follow-up area. For example, does DA bind directly to α1-ARs and stimulate Ca^2+^ independently from DRs? Or does DA induce a physical interaction between the bound DR and α1-AR, which then drives downstream Ca^2+^ mobilization? While radioligand binding studies indicate that non-specific interaction of DA with α1-AR only occurs at sub-millimolar concentrations (Proudman and Baker, 2021; Steinberg and Bilezikian, 1982) (much higher than those used in the present experiments), D1 and α1AR colocalize on PFC dendrites and have been suggested to undergo co-trafficking (Mitrano et al., 2014). In addition, mounting evidence from co-immunoprecipitation, BRET/FRET sensors and proximity-ligation assays support the idea that DRs can form functional heteromeric complexes with ARs and other class A GPCRs (Azdad et al., 2009; Bonaventura et al., 2014; González et al., 2012; Kolasa et al., 2018; Lee et al., 2004; Moreno et al., 2014; Navarro et al., 2018; Pelassa et al., 2019; Rebois et al., 2012; Trifilieff et al., 2011; Valle-León et al., 2021; Zhu et al., 2020), although evidence against the existence of DR heteromers *in vivo* also exists (Frederick et al., 2015).

Many clinical drugs used to treat psychiatric disorders such as depression, anxiety, ADHD, and schizophrenia target multiple monoamine systems, and can have severe side effects and low remission rates (Stanford and Heal, 2019). For instance, astrocytes have been linked to ADHD (Nagai et al., 2019), and methylphenidate—a first-line therapy for ADHD—increases both DA and NE concentration by blocking DAT and NET (Berridge et al., 2006). Our results here highlight open questions about treatments like these: are both neuromodulators needed for the therapeutic outcomes of methylphenidate, and are they both involved in its adverse effects? Do DA and NE act differently on neurons or non-neuronal cells? Are both DRs and ARs required to transduce DA signals in astrocytes, and if so would drugs that specifically target this interaction achieve better outcomes and minimize side effects? Clarifying the nature of the interaction between dopamine and ARs will be key for understanding the implications of drug treatment in conditions where both catecholamine systems are involved.

### Prefrontal DA, astrocytes and ATP

Extracellular ATP is an important regulator of brain circuits that can be released by astrocytes in other brain areas in response to DA (Corkrum et al., 2020) and other neurotransmitters (Gordon et al., 2005; Lalo et al., 2014; Pougnet et al., 2014) and lead to a generalized depression of synaptic transmission (Corkrum et al., 2020; Martin-Fernandez et al., 2017; Pascual et al., 2005; Zhang et al., 2003). PFC astrocytes are capable of ATP release (Cao et al., 2013), and, consistent with this, our experiments show regulation of extracellular ATP in response to DA (Fig 7). In our experimental conditions, the mobilization of ATP is delayed relative to DA, which could potentially tie this function of astrocytes to sustained activity in PFC during delay periods of working memory tasks. DA actions on both pyramidal cells and interneurons influence recurrent activity in PFC microcircuits (Goldman-Rakic, 1995), as well as excitatory/inhibitory balance (Gao and Goldman-Rakic, 2003; Gao et al., 2003), which are fundamental for the generation and maintenance of sustained activity. Because ATP depresses transmission, DA may favor suppression of PFC activity through astrocyte-derived ATP. Our ATP imaging shows that DA acting on PFC astrocytes induces spatially restricted patterns of extracellular ATP, suggesting that astrocytes could depress activity of specific PFC synapses located within their territories. Cell-type specific activity in PFC occurs during specific cognitive tasks (Kamigaki and Dan, 2017; Kim et al., 2016; Pinto and Dan, 2015), and cell-type specificity in astrocyte-neuron interactions has been reported in other brain areas (Martín et al., 2015; Martin-Fernandez et al., 2017), so future experiments could explore whether PFC astrocytes regulate ATP at defined pyramidal or interneuron subtypes, or subsets of synapses, to coordinate specific PFC microcircuits and influence cognitive function.

### Conclusion

Overall, the present work contributes to the fields of PFC, DA, and astrocyte research. Our data indicate that prefrontal astrocytes have distinct features compared to sensory cortex astrocytes, and respond to DA with non-canonical Ca^2+^ mobilizations that occur via α1-AR. Future work will address how PFC DA acting on astrocytes contributes to network dynamics and executive function, as well as the molecular basis and functional consequences of the crosstalk between DA and adrenergic receptors.

## Acknowledgements

The authors are grateful to members of the Poskanzer lab for helpful discussions, in particular Michelle Cahill for discussions about *ex vivo* analysis. We thank the Bender lab (UCSF) for reagents and assistance with PFC slicing, the Yuste lab (Columbia University) for assistance with reagents, Victoria Cheung in the Feinberg lab (UCSF) for help with initial fiber photometry recordings, Hajime Hirase (RIKEN, University of Copenhagen) for Pink Flamindo virus, and Jennifer Thompson for essential administrative support. K.P. is supported by NIH R01NS099254, NIH R01MH121446, NIH R21DA048497, and NSF CAREER 1942360. S.P. is supported by the EU under the Horizon 2020 Marie Skłodowska Curie Actions project ASTRALIS, GA 839561. M.R. was supported by the UCSF Genentech Fellowship. R.E. is a member of CONICET.

## Author Contributions

K.P and S.P. conceived and designed the study, and wrote the manuscript with input from the other authors. S.P. carried out *ex vivo* imaging and fiber photometry experiments and analyzed all data. S.Y. performed the *in vivo* 2P imaging experiments in PFC through GRIN lenses, *post hoc* immunostaining, and immunostaining/analysis of dopaminergic receptors. D.W. performed the fiber photometry experiments. C.T. performed the *ex vivo* GRAB_ATP_ experiments. M.R. carried out the *in vivo* 2P imaging experiments of V1. V.T. carried out immunostaining of *post hoc* tissue. Z.W. and Y.L. developed, tested, and provided GRAB_ATP_. R.E. provided critical support related to the RuBi-DA.

## Declaration of Interests

The authors declare no competing interests.

---

**Figure S1, related to Fig. 1.**
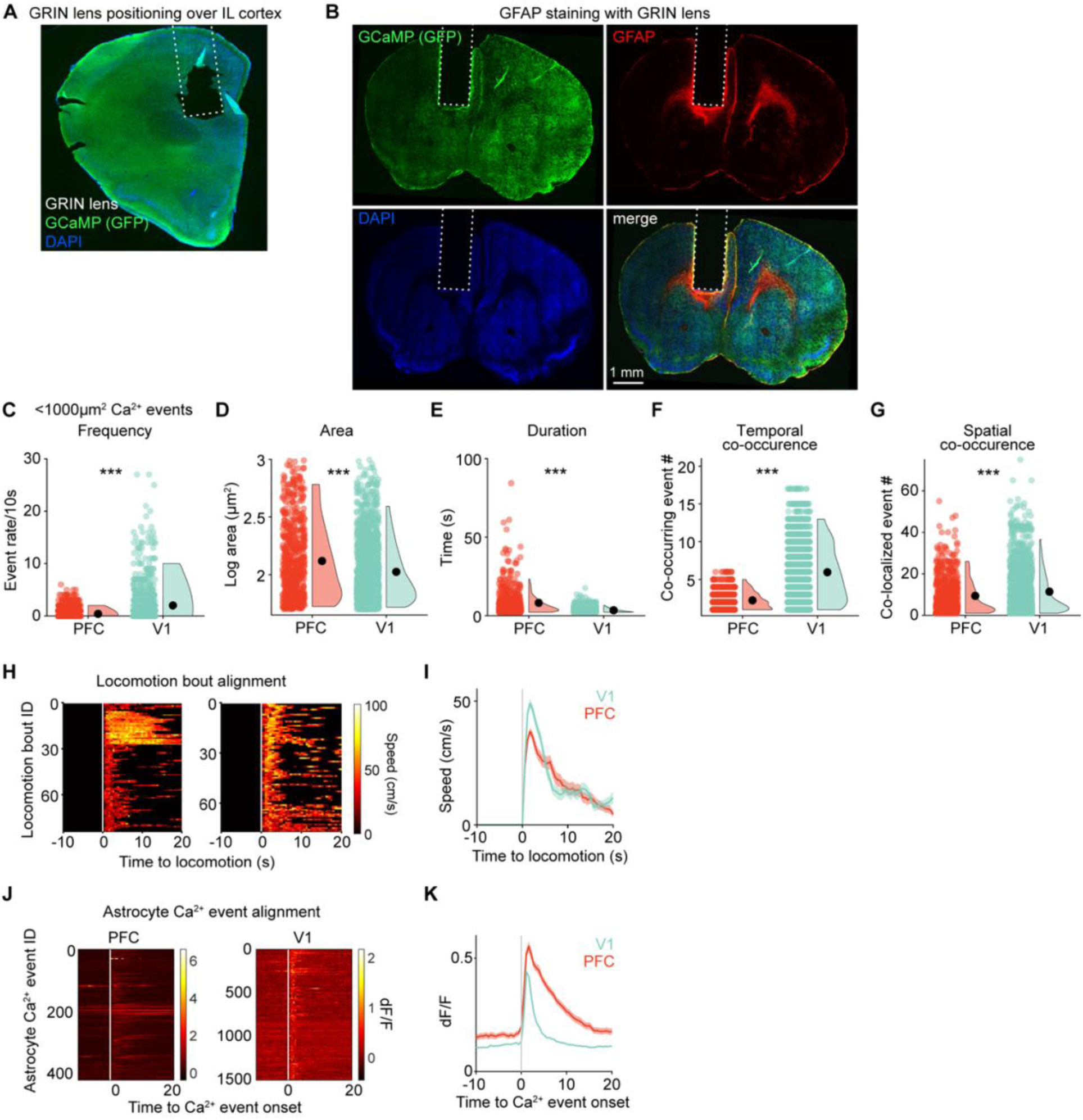
**(A)** GRIN lens implant location in PFC was confirmed by fixation and sectioning (∼2 mm anterior from Bregma), immunostaining to visualize astrocytic expression of Lck-GCaMP (green) and DAPI (blue) for nuclei. **(B)** GFAP (red), astrocytic Lck-GCaMP (green), and nuclei staining (DAPI, blue) were used to assess astrocyte reactivity around implanted GRIN lenses. **(C–G)** Small (<1000 µm^2^) astrocyte Ca^2+^-event features also vary between brain regions. Small events occur at lower rates in PFC (C), but are (D) larger and (E) longer than those in V1. Small events in PFC (F) co-occur with other events less than in V1, but (G) tend to repeat less at the same spatial location. Data shown as all bins/events (colored dots), 5^th^–95^th^ percentile distribution (violins), and mean ± sem (black dots and error bars). Wilcoxon rank-sum test; ***, *p* < 10^-5^. PFC: n = 1486 60-s bins, 593 events, 4 mice; V1: n = 780 60-s bins, 1561 events, 3 mice. **(H–I)** Animal speed (cm/s) aligned to start of locomotion bouts (t = 0), shown as heat map for all locomotion bouts (H) and average traces ± sem (I). **(J–K)** Astrocyte Ca^2+^ traces (dF/F) aligned to the onset of the AQuA-detected Ca^2+^ events (t = 0), shown as heatmaps for all events (J) and average traces ± sem (K).

**Figure S2, related to Fig. 2.**
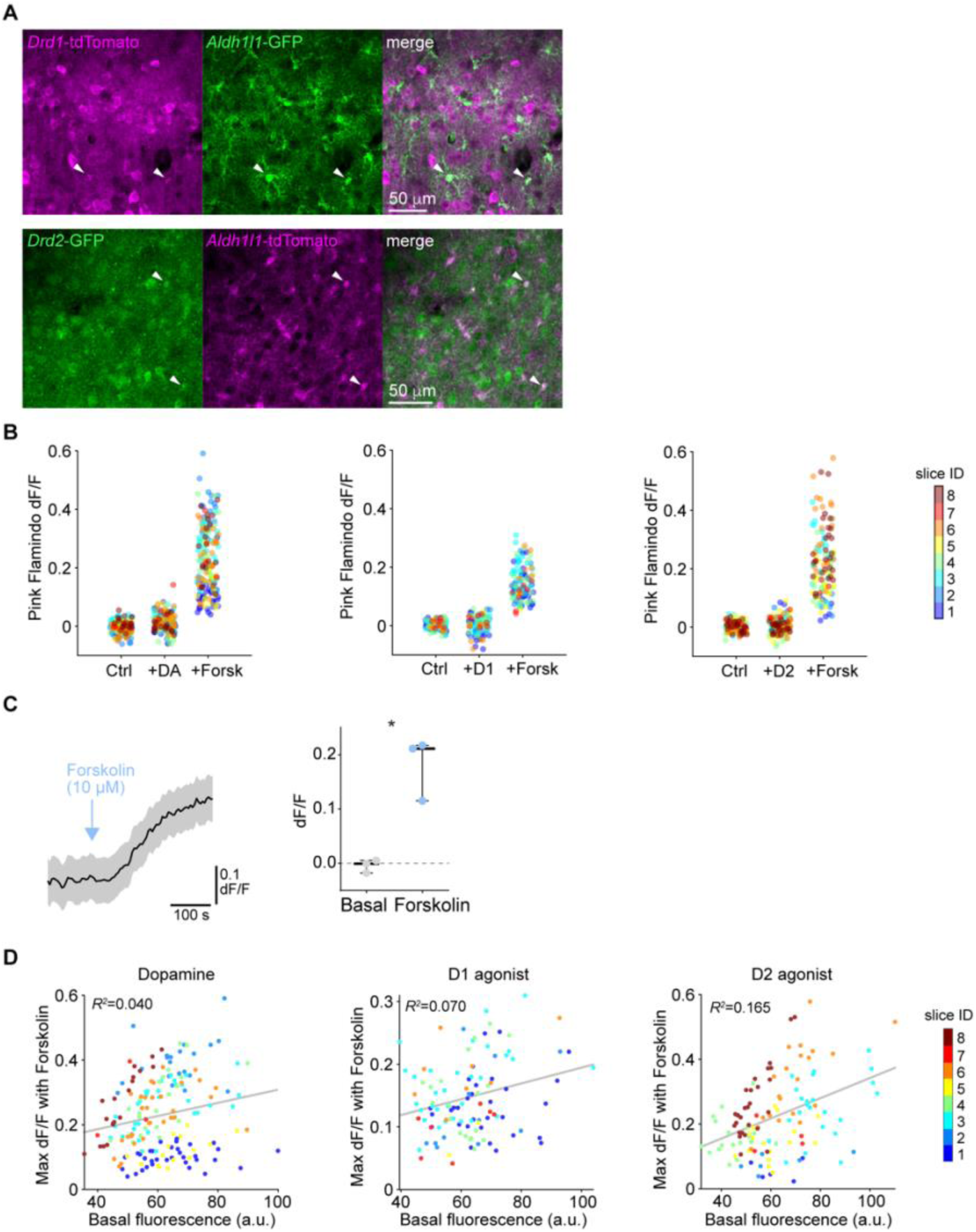
**(A)** Examples of high magnification (20x) micrographs from PFC of (top) *Drd1*-tdTomato x *Aldh1l1*-GFP and (bottom) *Drd2*-GFP x *Aldh1l1*-tdTomato mice to demonstrate colocalization and cell morphology. Arrowheads indicate cells co-expressing either receptor and Aldh1l1. **(B)** Fluorescence change from baseline for all analyzed cells, with colors indicating individual slices, to highlight no evident clustering of cells within individual slices. For the “+DA” condition, there is no evident separation of cells into two separate clusters, indicating that DA does not activate D1 (G_s_ pathway) and D2 receptors (G_i_ pathway) in different cells. Corresponding slice averages are shown in Fig 2G. **(C)** Adenylate cyclase activator Forskolin mobilizes cAMP in naïve slices, *i.e.* not treated with TTX and the drug cocktail to block neurons, at similar levels to those observed in treated slices (right; + Forskolin = 0.18 ± 0.03 dF/F). Data shown as (left) mean traces ± sem and (right) slice averages ± sem of whole-cell Pink Flamindo fluorescence (dF/F) before (basal) and after Forskolin. Paired t-test; *, *p* = 0.036; n = 58/144 cells, 3 slices and mice. Scale = 100 s, 0.1 dF/F. **(D)** Astrocytic response to maximal adenylate cyclase stimulation correlates weakly with cAMP reporter expression (Pink Flamindo basal fluorescence, arbitrary units). The observed variability in maximal cAMP concentrations in PFC astrocytes may reflect different signaling capabilities of individual astrocytes (*e.g.,* differential expression levels of AC subtypes, phosphodiesterases, *etc.*). Data are shown as cells (dots), colors indicate individual slice IDs. Pearson’s coefficients (*r^2^*) of linear correlation fits are indicated; p-values < 0.007.

**Figure S3, related to Fig. 3.**
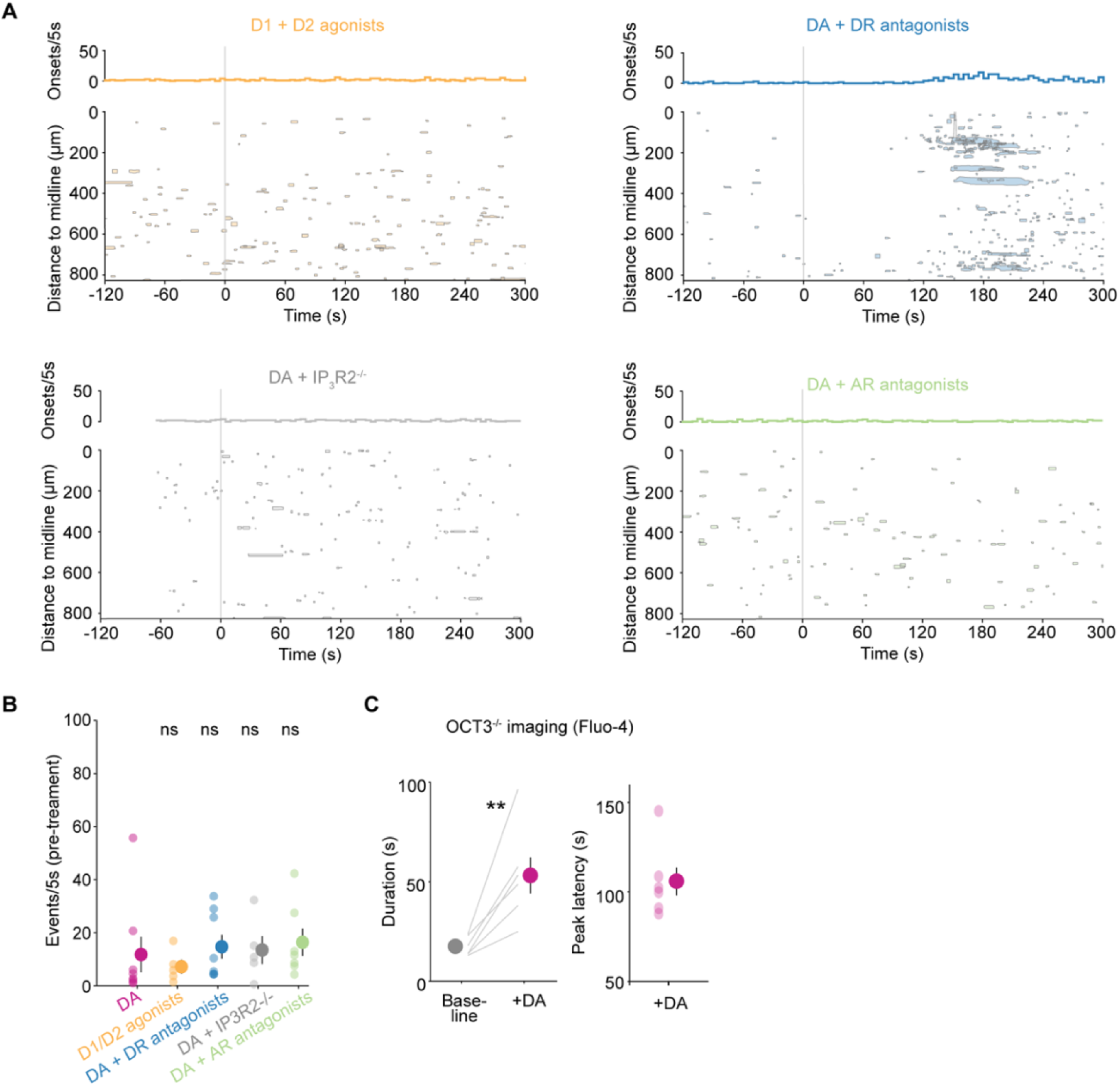
**(A)** Time course of all astrocyte Ca^2+^ events detected in PFC slices relative to D1/D2 agonists application (SKF38393/Quinpirole, top left), or relative to DA application in D1/D2 antagonists (SCH23390/Sulpiride, top right), in IP_3_R2 KO mice (bottom left), or in α-/β-AR antagonists (Phentolamine/Propranolol, bottom right). Rate of event onset (counts in 5-s bins) is displayed on top of each graph. Shaded areas represent approximate event size and mean y-position of the event over time. **(B)** No difference in pre-treatment event rate (count/5 s) for all slices and conditions shown in Fig. 2E. Event rate was calculated over a 60-s period before treatment with DA or D1/D2 agonists (as indicated by x-axis labels). Data shown as all slices (transparent dots) and corresponding mean ± sem (solid dot and error bar) for each condition): 11.8±6.7 (DA); 7.2±2.7 (D1/D2 ago.); 14.7±4.5 (DR antag.); 13.5±5.3 (IP_3_R2^-/-^); 16.4±5.2 (AR antag.). One-way Anova after Levene test; *p* = 0.829; n = 5–8 slices, 4–8 mice. **(C)** Average duration (left) and latency (right) of astrocytic somatic Ca^2+^ transients in slices from OCT3^-/-^ mice imaged using Fluo-4. Duration of Ca^2+^ transients is higher after DA compared to that of spontaneous Ca^2+^ activity (basal). In OCT3^-/-^ astrocytes, the latency to Ca^2+^ recruitment in response to DA is similar to that observed in wild-type mice (see Fig. 3D). Data shown as slice averages (lines or transparent dots) and corresponding means ± sem (solid dots and error bars). Duration (s): 17±2 (Baseline); 53±10 (DA). Peak latency: 106±8 s. Paired t-test after Anderson-Darling test; **, *p* = 0.009; n = 138 active cells, 6 slices, 3 mice.

**Figure S4, related to Fig. 4.**
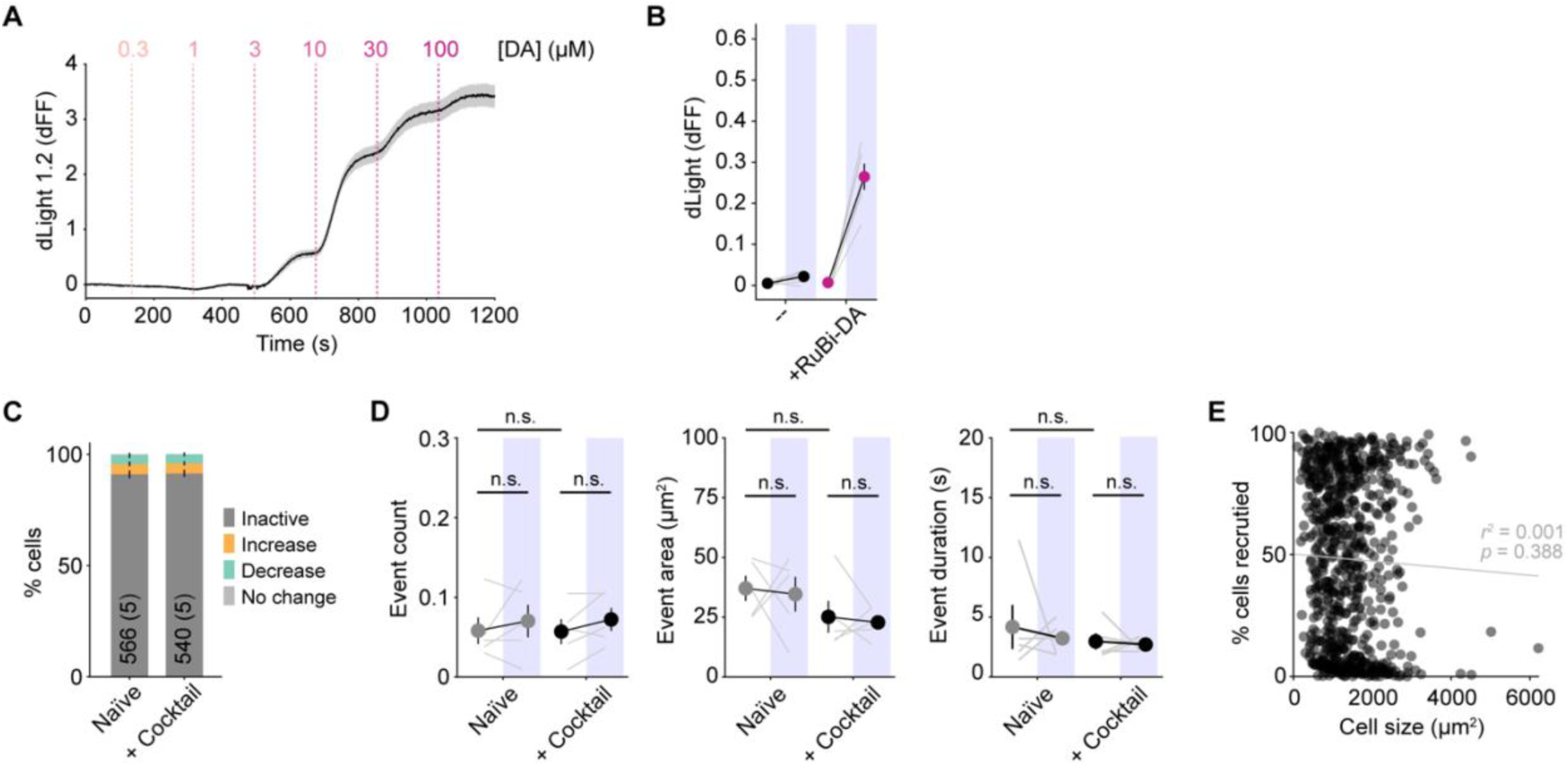
**(A)** Representative dLight dose-response experiment, showing time course of dLight fluorescence in slices challenged with increasing concentrations of DA (0.3 to 100 μM, in 1-log steps; times of application indicated by dotted lines and corresponding concentrations shown on top). Data shown as mean ± sem of different regions-of-interest selected across the imaging field. **(B)** dLight1.2 fluorescence before and after uncaging (across 30s, shaded blue boxes) in PFC slices, in absence (--, black dots) or presence of RuBi-DA (magenta). Data from Fig. 4D, shown as slices (grey lines) and corresponding mean ± sem (black lines, solid dots and error bars): 0.005±0.002, 0.022±0.005 (--); 0.007±0.001, 0.265±0.032 (+RuBi-DA). **(C)** Astrocytes were largely inactive (no Ca^2+^ events) before (naïve) and after addition of TTX and the multidrug cocktail (+ cocktail) used in all experiments in Fig. 4F–L. Percentage of cells increasing, maintaining, or decreasing their activity after the uncaging light pulse were low and similar between conditions, indicating no effect of light stimulation or drugs. Data shown as mean ± sem; n = 540–566 cells, 5 slices, 5 mice. **(D)** Ca^2+^ event features (number, area and duration) in active cells in (C) were unchanged after addition of TTX and multidrug cocktail. There was similarly no effect of the uncaging light (shaded blue boxes) on event features in naïve slices and those treated with the cocktail. Data shown as slice means (grey lines) and mean ± sem (dots and error bars). After checking normality, the Wilcoxon rank sum test was used to compare event features before uncaging between naïve and cocktail, and the paired t-test used to compare event features pre- versus post-uncaging. Naïve: n = 52/566 cells, 5 slices and mice. Cocktail: n = 47/540 cells, 5 slices, 5 mice. **(E)** The maximum cell area recruited by fast release of DA is independent of cell size, indicating that our cell delineation method does not affect this measurement. *R^2^*, Pearson’s coefficient, and *p*, p-value of linear correlation fit (grey line). N = 1118 cells, 11 slices, 8 mice.

**Figure S5, related to Fig. 5.**
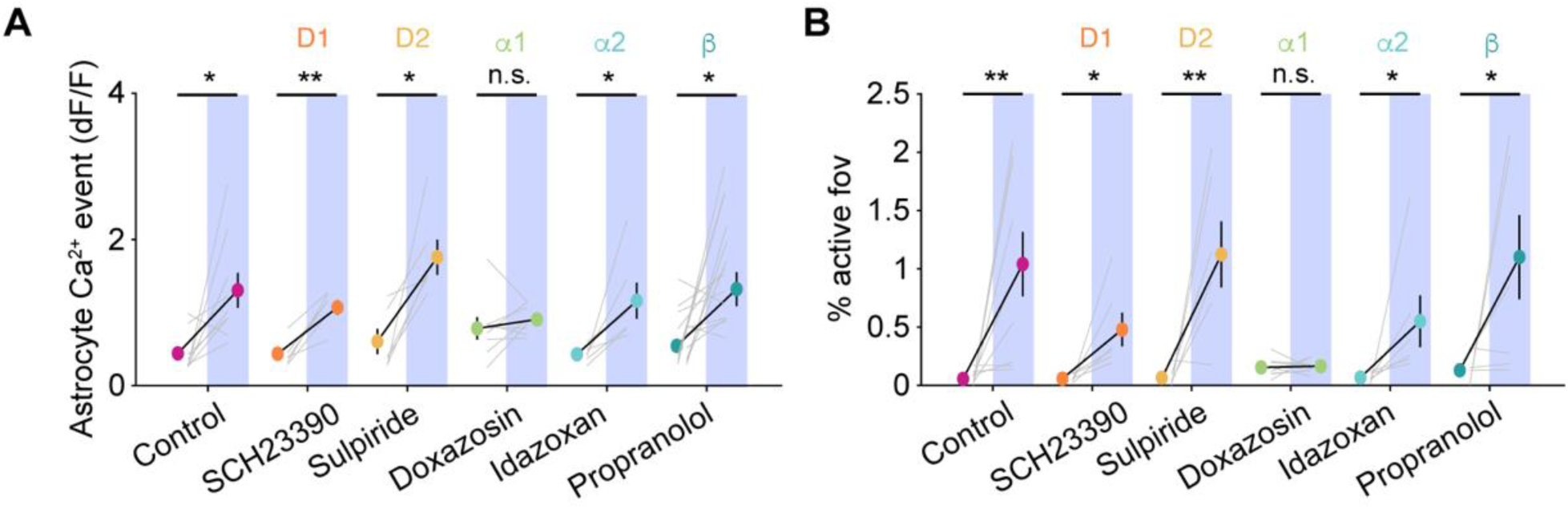
**(A–B)** Additional quantification of experiments in Fig. 5B yield comparable results to z-scored traces of AQuA Ca^2+^-events (Fig. 5C). **(A)** dF/F traces of AQuA Ca^2+^-events. **(B)** Percent of active imaging field recruited over time. Data shown as 30-s means of slice Ca^2+^ immediately before or after RuBi-DA uncaging (shaded blue boxes) in presence of different receptor inhibitors as indicated, for slice averages (grey lines) and corresponding mean ± sem (black lines, solid dots and error bars). One or two-tailed paired t-test or Wilcoxon signed rank test after checking normality with the Anderson-Darling test to compare pre- to post-uncaging values; *, *p* < 0.05, **, *p* < 0.01; n = 6–9 slices, 5–9 mice.

**Figure S6: Related to Fig. 6.**
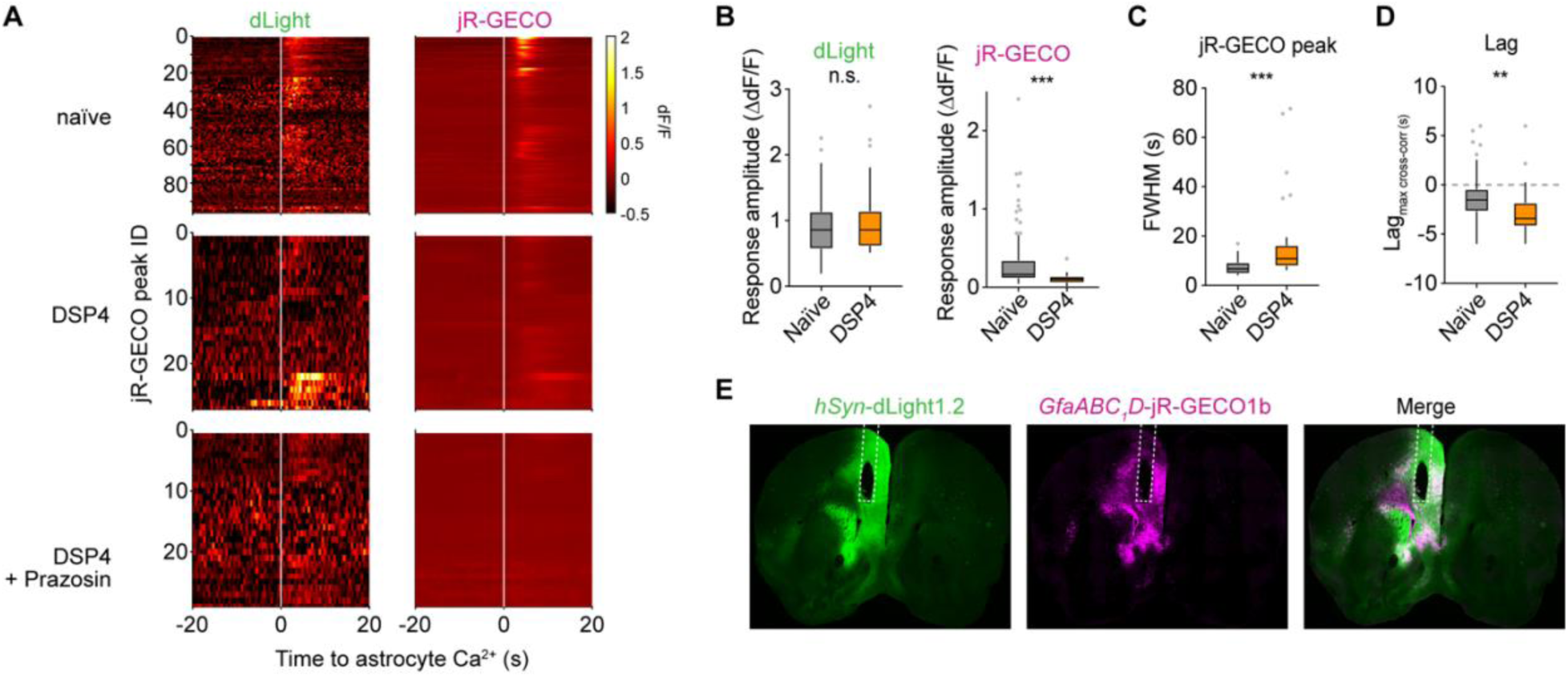
**(A)** Fiber photometry recordings of dLight (left) and astrocytic jR-GECO (right) aligned to the onset of astrocyte Ca^2+^ transients evoked by aversive stimulation (t = 0), shown as heat map of dF/F for all jR-GECO peaks detected and conditions (top: naïve; middle: DSP4; bottom: DSP4 and Prazosin). **(B)** Amplitude of DA transients (left, dLight) in PFC (naïve: 0.89±0.04 dF/F) was not significantly affected by ablation of LC fibers (DSP4: 1.02±0.11 dF/F), whereas astrocyte Ca^2+^ transients (right, jR-GECO) were significantly lower with decreased NE input (naïve: 0.31±0.04 dF/F; DSP4: 0.11±0.01 dF/F), indicating that the integrity of the NE system is important for astrocyte function in PFC. Data shown as Tukey boxplots. Wilcoxon rank sum test; ***, *p* < 0.001; n = 27–96 transients, 4–9 mice. **(C)** Astrocyte Ca^2+^ transients (jR-GECO peaks) were longer after decreased LC input to the PFC (naïve: 7.3±0.3 s; DSP4: 18.3±3.5 s). Data shown as Tukey boxplots. Wilcoxon rank sum test; ***, *p* < 0.001; n = 27–96 transients, 4–9 mice. **(D)** Lag of maximum cross-correlation between dLight and jR-GECO was significantly higher after decreased LC input to the PFC (naïve: -1.4±0.2 s; DSP4: -2.6±0.5 s). Data shown as Tukey boxplots. Wilcoxon rank sum test; **, *p* = 0.004; n = 27–96 transients, 4–9 mice. **(E)** Optic fiber implant location in PFC was confirmed by immunostaining ∼2 mm rostral from Bregma to visualize expression of dLight (green) and jR-GECO1b in astrocytes (magenta), but note that implant location was determined initially by the stereotax coordinates.

## Supplemental Movies

**Movie 1: PFC astrocytes *in vivo* display cell-restricted Ca^2+^ activity unlinked to animal locomotion.**

Ca^2+^ events in PFC (left) and V1 (right) astrocytes, with their relationship to animal locomotion. Top: raw Lck-GCaMP6f movies with overlaid AQuA-detected Ca^2+^ events (colors are individual events). Bottom: time course of astrocyte activity shown as population-level Ca^2+^ traces with corresponding animal locomotion speed below. Images acquired at 2 Hz, playback speed 15 fps.

**Movie 2: Robust astrocyte Ca^2+^ response to DA stimulation of PFC slices.**

Bath-application of DA (10 μM) in PFC slices expressing GCaMP6f in astrocytes induces a robust, but delayed Ca^2+^ mobilization. Neuronal action potentials and neuron-to-astrocyte communication are blocked with TTX and a drug cocktail (see Methods). Top: raw movie; bottom: AQuA-detected Ca^2+^ events. Time stamps relative to DA application.

**Movie 3: DA uncaging induces fast, transient Ca^2+^ activation in PFC astrocytes.**

RuBi-DA (right) uncaged at t=0 on GCaMP6f-expressing astrocytes in a PFC slice, inducing a robust Ca^2+^ response within seconds. In the same slice, the light stimulation protocol in the absence of RuBi-DA (control, left) has no effect. Top: raw movies; bottom: overlaid AQuA-detected Ca^2+^ events (colors are individual events). Time indicated in seconds from uncaging.

**Movie 4: DA acts on PFC astrocytes via α1-adrenergic receptors.**

Representative experiments in PFC slices expressing GCaMP6f in astrocytes, in which RuBi-DA is uncaged at t = 0 s in presence of different DR and AR antagonists. Labels indicate drugs (Control: no antagonist; D1: SCH23390 10 μM; D2: Sulpiride 0.5 μM; α1: Doxazosin 10 μM; α2: Idazoxan 10 μM; β: Propranolol 10 μM). Uncaging light pulses appear as white stripes due to detector saturation. Only Doxazosin application blocks uncaging-induced astrocyte activation.

## Methods

### Animals

Experiments were carried out using young adult for *ex vivo* or adult mice for *in vivo* experiments, in accordance with protocols approved by the University of California, San Francisco Institutional Animal Care and Use Committee (IACUC). Animals were housed in a 12:12 light-dark cycle with food and water provided *ad libitum*. Male and female mice were used whenever available. For *in vivo* experiments following surgery, all animals were singly housed to protect implants and given additional enrichment. Transgenic mice used in this study were Lck-GCaMP6f^fl/fl^ mice (Srinivasan et al., 2016) and *Aldh1l1*-Cre/ERT2 mice (Srinivasan et al., 2016) from the Khakh lab (UCLA, USA), *Drd1a*-tdTomato (Shuen et al., 2008) and *Drd2*-EGFP (Gong et al., 2003) from the Bender lab (UCSF, USA), *Aldh1l1*-EGFP and *Aldh1l1*-tdTomato (Gong et al., 2003) from JAX (USA), *Itpr2*-deficient mice (IP_3_R2^-/-^) (Li et al., 2005) from Dr. Katsuhiko Mikoshiba (RIKEN, Japan) and *Slc22a3*-deficient mice (OCT3^-/-^) (Zwart et al., 2001) from the Irannejad lab (UCSF, USA).

### Surgical procedures

For viral expression in *ex vivo* experiments, neonatal mice (P0–4) on C57Bl/6 or Swiss background were anesthetized on ice for 2 min before injecting viral vectors (*AAV5-GfaABC_1_D-GCaMP6f* [1.4–5.42e^13^; all titers in GC/ml], *AAV9-hGfap-pinkFlamindo* [6.6e^13^], *AAV9-hSyn-NE2.1* [5.72e^13^], *AAV9-CAG-dLight1.2* [9.5e^15^], or *AAV9-hSyn-ATP1.0* [4.89e^13^]). Pups were placed on a digital stereotax and coordinates were zeroed at the middle point along the line connecting the eyeballs. Two injection sites over PFC were chosen at 0.25–0.34 mm lateral, and 1 and 1.4 mm caudal. At each injection site, 30–100nl of virus were injected at a rate of 3–5nl/s at two depths (0.7–0.85, and 0.9–1 mm ventral) using a microsyringe pump (UMP-3, World Precision Instruments).

For *in vivo* 2P imaging, we expressed Lck-GCaMP in astrocytes of adult mice (P50–130), either by crossing Lck-GCaMP6f^fl/fl^ mice to *Aldh1l1*-Cre/ERT2 (Srinivasan et al., 2016) and treating them with tamoxifen (0.1 mg/kg, i.p., for 5 consecutive days, 4–6 weeks before imaging), or via viral vectors (see below) in C57Bl/6 mice. For fiber photometry, we expressed dLight and astrocytic jR-GECO1b via viral vectors (see below) in C57Bl/6 mice (P60–90). Before surgical procedures, adult mice were administered dexamethasone (5 mg/kg, s.c.) and anesthetized with isoflurane, and a 1 or 3-mm diameter craniotomy was created over PFC (+1.7–1.8 mm rostral, +0.5 mm lateral from Bregma) or visual cortex (−3.5 mm caudal, +1.2 mm lateral from Bregma). Viral vectors (*AAV5-GfaABC_1_D -Lck-GCaMP6f-SV40* [1.4–5.42e^13^], *AAV5-hSyn-dLight1.2* [4e^12^], *AAV9-GfaABC_1_D-Lck-jRGECO1b* [2.24e^14^]) were delivered using a microsyringe pump (100–600 nl, 30–60 nl/min) before implanting optical devices. For 2P imaging in PFC, after careful removal of meninges, a GRIN lens (1-mm diameter, 4.38-mm length, WDA 100, 860 nm, Inscopix) was slowly lowered to -2.4 mm ventral; for 2P imaging in V1, a cranial window was placed above the tissue; a custom-made titanium headplate was then attached to the skull. For fiber photometry in PFC, a fiber optic cannula (Mono Fiberoptic Cannula, 400μm core, 430nm, 0.48 NA, 2.8mm length, Doric Lenses) was implanted at the same depth as GRIN lenses. All imaging devices were secured in place using dental cement (C&B Metabond, Parkell). Post-operative care included administration of 0.05 mg/kg buprenorphine and 5 mg/kg carprofen. Mice were allowed a minimum of 14 days to recover, then habituated to head-fixation on a circular treadmill or to fiber optic coupling in a freely moving arena prior to experiments.

### *In vivo* 2P imaging and locomotion

2P imaging experiments were carried out on an upright microscope (Bruker Ultima IV) equipped with a Ti:Sa laser (MaiTai, SpectraPhysics). The laser beam intensity was modulated using a Pockels cell (Conoptics) and scanned with linear galvanometers. Images were acquired with a 16×, 0.8 N.A. (Nikon) or a 20×, 1.0 N.A. (XLUMPLFLN-W, Olympus) water-immersion objective via photomultiplier tubes (Hamamatsu) using PrairieView (Bruker) software. For GCaMP imaging, 950 nm excitation and a 515/30 emission filter were used. Mice were head-fixed on a circular treadmill and Ca^2+^ activity was recorded at ∼1.7 Hz frame rate from putative PFC or V1 cortex, at 512×512 pixels and ∼0.6 μm/px resolution. Locomotion speed was monitored using an optoswitch (50mA, 2V; OPB800L55, TT Electronics, Newark) connected to a microcontroller board (Arduino Uno R3, Arduino) and acquired at 1KHz simultaneously with 2P imaging using PrairieView.

### *Ex vivo* 2P imaging and uncaging

Coronal, acute PFC slices (300-μm thick) from P27–P54 mice were cut with a vibratome (VT 1200, Leica) in ice-cold cutting solution containing (in mM) 27 NaHCO_3_, 1.5 NaH_2_PO_4_, 222 sucrose, 2.6 KCl, 2 MgSO_4_, 2 CaCl_2_. Slices were transferred to pre-heated, continuously aerated (95% O_2_/5% CO_2_) standard artificial cerebrospinal fluid (ACSF) containing (in mM) 123 NaCl, 26 NaHCO_3_, 1 NaH_2_PO_4_, 10 dextrose, 3 KCl, 2 MgSO_4_, 2 CaCl_2_. Younger mice were sliced in the same solutions for dLight (P18–28) and GRAB_NE_ (P24–35) experiments, and one P19 IP_3_R2^-/-^ experiment (otherwise P31–36). Slices were kept at room temperature until imaging, and experiments performed at 37°C. To block neuronal action potentials and neuron-to-astrocyte-communication during imaging, at least 10 min before experiments recirculating standard ACSF was switched to a multi-drug cocktail mix, containing (in µM) 1 TTX, 100 LY341495, 1 CGP 55845, 2 AM251, 1 CGS 15943, 100 PPADS, 5 Ipratropium, unless otherwise stated.

Slice recordings were done in coronal sections above medial prefrontal cortex, and the imaging area was ∼ 0.6–0.8 mm x 0.8 mm over prelimbic and infralimbic areas, with the top part of the imaging area corresponding to the midline, thus spanning all cortical layers. Images were acquired from putative PL-IL cortex in PFC slices at a minimum depth of 50 μm, using the same setup as for *in vivo* 2P imaging or a custom-made upright microscope and ScanImage software, at 1.42–1.53 Hz frame rate, 512×512 pixels and 1.04–1.61 μm/px resolution. Fluorophores were excited at (in nm) 950–980 (GCaMP), 1040 (Pink Flamindo), 980 (dLight), 920 (GRAB_NE_ and GRAB_ATP_). Emission was collected with a 515/30 or 525/50 filter for green and a 605/15 or 600/40 filter for red fluorophores. For bath-application experiments, a 5-min baseline was recorded to monitor spontaneous activity, after which neuromodulators were added along with a fluorescent dye (AlexaFluor 594 Hydrazide) to assess the time at which drugs reached the imaging field (except for Pink Flamindo due to spectral overlap).

For RuBi-DA uncaging, a fiber optic cannula (400-µm core, 0.39 NA; CFM14L10, ThorLabs) was coupled to a compatible fiber optic (M79L005, ThorLabs) and a blue LED (470 nm; M470F3, ThorLabs), and placed adjacent to the imaging field using a micromanipulator (MX160R, Siskiyou). Illumination (3 pulses, 100-ms duration, 50-ms intervals) was triggered using the imaging software (PrairieView, Bruker) connected to the LED-driver cube (LEDD1B, ThorLabs). Light power was 2–4 mW.

### Fiber photometry recordings

FP experiments were carried out using an RZ10 fiber photometry processor equipped with Lux integrated 405, 465, and 560-nm LEDs and photosensors (Tucker-Davis Technologies). Animals implanted for FP were placed in a freely moving arena in which the mouse was able to move in all directions, after coupling to low autofluorescence fiberoptic patchcords (400-μm core, 0.57 NA; Doric Lenses) connected to photosensors through a rotary joint (Doric Lenses). FP fluorescence signals were recorded for 10 minutes, during which tail lifts were performed every minute. For a tail lift stimulation, the experimenter held and lifted the tail of the animal until its hind paws disconnected from the ground; after that the tail was released. With this experimental paradigm, no pain or harm is caused to the animal. After baseline recordings, animals were treated with DSP4 (50 mg/kg, i.p., 2 injections 2 days apart) and recorded again 4 days after the first DSP4 administration. Following DSP4 recordings, animals were injected with Prazosin (5 mg/kg, i.p.) and recorded again 20 minutes later.

### Immunohistochemistry

Mice were intracardially perfused with 4% PFA, brains were then collected, immersed in 4% PFA overnight at 4°C and switched to 30% sucrose for two days before being frozen on dry ice and stored at - 80°C. Brains were sliced coronally (40-μm thick) on a cryostat, and slices stored in cryoprotectant at - 20°C until staining. Slices were washed 3x in PBS for 5 min, then permeabilized for 30 min with 0.01% TritonX in PBS. Slices were next washed with 10% NGS (Invitrogen) for 1 h and incubated overnight with primary antibodies at 4°C in 2% NGS. Slices were next rinsed 3x in PBS before incubating for 2 h at room temperature with secondary antibodies, then washed 3x in PBS for 5 min before slide-mounting and coverslipping using Fluoromount with DAPI.

To stain for EGFP and tdTomato in D1, D2 and Aldh1l1 colocalization experiments, primary antibodies used were rat α-mCherry (1:1000, Thermo Fisher Scientific) and chicken α-GFP (1:3000, Aves Lab) in 2% NGS. Secondary antibodies used were goat α-rat Alexa Fluor 555 (1:1000) and goat α-chicken Alexa Fluor 488 (1:1000). To estimate colocalization, 3 slices/mouse were chosen at +1.8, +1.7 and +1.6 mm from bregma, and tiled z-stack images were acquired on a spinning disk confocal (Zeiss) at PFC spanning cortical layers 1–6. Colocalization counts of tdTomato^+^ and EGFP^+^ cells were performed using Cell Counter in Fiji (ImageJ).

To stain brain tissue from GRIN lens experiments, primary antibodies used were rat α-GFAP (1:1000, Thermo Fisher Scientific) and chicken α-GFP (1:3000, Aves Lab) for Lck-GCaMP. To stain for dLight and jR-GECO1b in sections from fiber photometry experiments, primary antibodies used were rat α-mCherry (1:1000, Thermo Fisher Scientific) and chicken α-GFP (1:3000, Abcam). Secondary antibodies used were goat α-rat Alexa Fluor 555 (1:1000, Thermo Fisher Scientific) and goat α-chicken Alexa Fluor 488 (1:1000, Thermo Fisher Scientific). For NET staining, sections were incubated for 1 h with a secondary mouse block (AffiniPure Fab Fragment IgG, 30 μg/ml, Jackson ImmunoResearch) before primary antibody mouse α-NET (1:100, MAb Technologies), and secondary antibody goat α-mouse Alexa Fluor 555 (1:1000, Thermo Fisher Scientific). Z-stacks or whole-brain images were acquired at 40x or 2x using a Keyence BZ-X800 fluorescence microscope and stitched with Keyence Analysis Software.

### 2P image and data analysis

When necessary, videos were preprocessed by registering images using the ImageJ plugin MoCo (Dubbs et al., 2016). Cell maps for Pink Flamindo, GRAB_ATP_, and uncaging experiments were drawn using the interactive wand segmentation tool (SCF-MPI-CBG plugin).

#### AQuA

Ca^2+^ and ATP 2P image sequences were analyzed using AQuA software (Wang et al., 2019) and custom MATLAB (Mathworks) code. Signal detection thresholds were adjusted for each video to account for differences in noise levels after manually checking for accurate AQuA-detection. Events were thresholded post-detection at 25 μm^2^ and 2 s for *ex vivo*, or 50 μm^2^ and 2 s for *in vivo* Ca^2+^ imaging, and at 50 μm^2^ and 2.5 s for GRAB_ATP_ imaging. Event count was quantified using the onset of each event as detected by AQuA. Area is defined as the footprint occupied by an event over its entire lifetime. Number of co-occurring events is calculated as the number of events co-existing temporally anywhere in the imaging field with a given event. Number of co-localized events is calculated as the number of events having comparable size (0.5–2x) and overlapping spatially with a given event.

#### *In vivo* 2P Ca^2+^ imaging and locomotion analysis

For locomotion-aligned astrocyte Ca^2+^ analysis, only locomotion bouts longer than 2 s and starting more than 10 s after the previous locomotion bout ended were considered (Fig S1H). Population-wide mean Ca^2+^ traces (Fig 1I) were obtained by normalizing the fluorescence of each AQuA-detected event as (F-F_min_)/(F_max_-F_min_) and then averaging across events. For max Ca^2+^ (Fig 1K), changes in normalized fluorescence were thresholded at 0.1 to exclude noise. For astrocyte Ca^2+^-aligned locomotion analysis (Fig 1L), astrocyte Ca^2+^ event dF/F was used and all locomotion bouts were considered. Locomotion speed was calculated as cm/s.

#### Bath-applied DA analysis

For bathed-DA experiments, Ca^2+^ event rate was calculated as counts of AQuA-event onsets in 5-s bins (Ca^2+^), and events for the post-treatment condition (Fig 3E) were analyzed over a 30-s window centered at 90-s post drug or at the timepoint when the event rate exceeds baseline (6 STD of event rate at baseline). We then calculated the peak onset as the last local minimum before the peak, and—to overcome false positives due to noise—we constrained the local minima to be below the 6 STD threshold for peak detection. At baseline (Fig S3B) we used a 60-s window to account for low number of spontaneous events. For GRAB_ATP_, events were analyzed over 300-s windows, immediately before (basal) and 90-s post drug. The 300-s window (for the post-drug condition) was started 90-s after the delivery of the drug since we wanted to probe ATP events that would follow DA-induced Ca^2+^ events, which—in bath application experiments (Fig. 3)—started ∼90 s after drug delivery.

#### Single-cell *ex vivo* uncaging analysis

Classification of cell activity around uncaging was done based on counts of AQuA-event onsets in the 60-s before versus 60-s after uncaging (t=0). Event features (count, area, duration) were averaged by cell and slice using the same temporal windows. Traces for the % of cell area active were obtained as the overall number of pixels/frame occupied by AQuA-detected events within an individual astrocyte (cell territories were defined by cell maps, see above). Traces were analyzed with custom-written code in MATLAB to find peak times, amplitudes (max % cell surface active) and duration (FWHM). Latency to peak onset after uncaging (delay) was obtained as the first timepoint above threshold (6 STD of the pre-uncaging activity).

#### ROI-based analysis

Pink Flamindo, GRAB_NE_ and dLight videos were analyzed using ROI-based approaches in ImageJ. Changes in fluorescence intensity were calculated as (F-F_0_)/F_0_ (dF/F), where F_0_ is the average intensity of the first 20–30 frames. For GRAB_NE_, dF/F values were extracted as 20-s means at 50 s before (pre-drug) or 340 s after compound addition. Fluo-4 videos from OCT3 KO experiments were analyzed using CalTracer 3 Beta (Poskanzer and Yuste, 2016), dF/F traces extracted from the automatically detected cell somata, and identified peaks checked manually for accurate detection before extracting duration and latency. Traces dF/Fs were then obtained as 5-s means at 100 s before or after DA addition based on average peak latencies. Data for the dLight dose-response curves were fit to a Hill equation (y = a + (b-a)/(1+10^((c-x)*d))), and DA concentrations released by RuBi-DA uncaging were extrapolated from the obtained fit function based on changes in dLight fluorescence after uncaging.

#### Pink Flamindo analysis

For Pink Flamindo experiments, background fluorescence was subtracted from raw fluorescence traces. To identify steady-state increases or decreases in fluorescence, traces were smoothened using a moving average and then fit using a modified Boltzmann’s sigmoidal equation y=a+(b-a)/(1+exp((c-x)/d), where a is the bottom, b is the top, c is the inflection point and d is the slope, using a nonlinear least squares algorithm (Levenberg-Marquardt) in MATLAB. Fit constraints were (b-a)>noise, slope<10, and inflection point at x>0. Cells where the sigmoid fit of the trace in response to Forskolin did not converge were excluded from all previous analyses. Cells with high noise (>0.1 dF/F) or drift (when change in dF/F before drug application exceeded noise) were removed. Noise was calculated as 3 STD at baseline. Average dF/F values (Fig 2G) were then extracted as 20-s means at 40 s before (control) or 240 s after compound addition (drug/Forskolin) from original traces.

### Fiber photometry analysis

FP data were preprocessed by downsampling and subtraction of the isosbestic channel linear fit (as in https://www.tdt.com/docs/sdk/offline-data-analysis/offline-data-python/FibPhoEpocAveraging), detrended to correct bleaching, and dF/F calculated as above (F_0_ obtained at 0–15 s). Traces were then denoised using an IIR lowpass filter in MATLAB (cutoff frequency 1Hz, steepness 0.95). Transients in jR-GECO1b traces were detected using the ‘findpeaks’ function in MATLAB (applied over normalized traces, with minimum peak height and prominence set to 25% and at a minimum distance of 20 s, according to the timing of the tail lift stimulation protocol. All trials/animals were analyzed using the same parameters for peak detection.) Transient onsets were determined as the timepoints where the first derivative exceeded 1 STD. Then, dLight and jR-GECO1b traces were extracted in 40-s windows centered at onsets, and the cross-correlation function calculated from the extracted traces with a 6-s maximum lag to obtain the latency to maximum cross-correlation. Response amplitudes (Fig 6C,G) were calculated for each detected peak as change in dF/F between trace average before onset and trace maximum after onset.

### Statistics

To compare one group of data with a hypothesized mean value we used a one-sample t-test or sign test (Wilcoxon) as appropriate after a normality test. When comparing two unpaired groups, we used the two-sample, unpaired t-test or the Wilcoxon rank sum test (Mann-Whitney) as appropriate after a normality test. When comparing two paired groups of data, we used the paired t-test or the Wilcoxon signed rank test after checking for normality on the difference between groups. Normality was checked using the Anderson-Darling test. When comparing treatments for three or more groups (Fig 3E,G) we used one-way Anova or the Kruskal-Wallis test after testing for equal variances using the Levene test (quadratic). For Pink Flamindo data (Fig 2G), we used the non-parametric Friedman test for paired data after the Levene test to compare within conditions (control, drug, Forskolin), and one-way Anova or Kruskal-Wallis test after the Levene test to compare across treatments. Multiple comparisons in Fig. 5 were not corrected *post hoc* to minimize type II errors (i.e., to avoid increasing the rate of false negatives; in a false negative, pre- to post-uncaging values would be erroneously considered non-significantly different from each other). All statistical tests are two-tailed unless otherwise stated in the figure legend (Fig. S5B).

Statistical significance for time-series data was computed using the shuffle test with custom-written code in MATLAB. Data pairs were selected as a reference value (trace mean from t<0 or the entire time window analyzed) and a given timepoint in the time-series (t>0 or all timepoints in the window). Data from the two groups were pair-wise shuffled for 10000 repetitions to calculate the difference between the two populations, the significance level α for rejecting H_0_ was set to 0.01, and Bonferroni correction was applied to account for multiple comparisons.

## Notes

### Competing Interest Statement

The authors have declared no competing interest.

